# Nucleoplasmic Lamin C Rapidly Accumulates at Sites of Nuclear Envelope Rupture with BAF and cGAS

**DOI:** 10.1101/2022.01.05.475028

**Authors:** Yohei Kono, Stephen A. Adam, Karen L. Reddy, Yixian Zheng, Ohad Medalia, Robert D. Goldman, Hiroshi Kimura, Takeshi Shimi

**Affiliations:** Cell Biology Center, Institute of Innovative Research, Tokyo Institute of Technology, Yokohama, Japan.; Department of Cell and Developmental Biology, Feinberg School of Medicine, Northwestern University, Chicago, IL.; Department of Biological Chemistry, Johns Hopkins University, Baltimore, MD.; Department of Embryology, Carnegie Institution for Science, Baltimore, MD.; Department of Biochemistry, University of Zurich, Zurich, Switzerland.; World Research Hub Initiative, Institute of Innovative Research, Tokyo Institute of Technology, Yokohama, Japan.

## Abstract

In mammalian cell nuclei, the nuclear lamina (NL) underlies the nuclear envelope (NE) to maintain nuclear structure. The nuclear lamins, the major structural components of the NL, are involved in the protection against NE rupture induced by mechanical stress. However, the specific role of the lamins in repair of NE ruptures has not been fully determined. Our analyses using immunofluorescence and live-cell imaging revealed that lamin C but not the other lamin isoforms rapidly accumulated at sites of NE rupture induced by laser microirradiation in mouse embryonic fibroblasts. The immunoglobulin-like fold domain and the NLS were required for the recruitment from the nucleoplasm to the rupture sites with the Barrier-to-autointegration factor (BAF). The accumulation of nuclear BAF and cytoplasmic cyclic GMP-AMP synthase (cGAS) at the rupture sites was in part dependent on lamin A/C. These results suggest that nucleoplasmic lamin C, BAF and cGAS concertedly accumulate at sites of NE rupture for repair.

**Summary:** Kono et al. show the rapid recruitment of nucleoplasmic lamin C to sites of nuclear envelope rupture with Barrier-to-autointegration factor. Lamin A/C is also involved in nuclear DNA sensing with cytoplasmic cGAS at the ruptured sites.

## Introduction

The genomic DNA in a mammalian cell is folded into higher-order chromatin structures in the nucleus, which is separated from the cytoplasm by the nuclear envelope (NE). The NE is bounded by a double lipid bilayer comprising the inner nuclear membrane (INM) and the outer nuclear membrane (ONM). The nuclear lamina (NL) underlies the INM at its nucleoplasmic face where it interacts with heterochromatin to regulate the size, shape and stiffness of the nucleus (Lammerding et al., 2004; Levy and Heald, 2010; Shimi et al., 2008; Swift et al., 2013). The LINC complex (Linker of Nucleoskeleton and Cytoskeleton) mediates the physical interactions between the NL, the intermembrane space, the ONM and the major cytoskeletal networks to propagate signals from the cell surface to the nucleus by mechanotransduction (Crisp et al., 2006). Nuclear pore complexes (NPCs) penetrate the INM and ONM and associate with euchromatin to control macromolecular trafficking between the nucleus and cytoplasm (Tran and Wente, 2006) and are also connected to the cytoskeleton (Gray and Westrum, 1976; Mahamid et al., 2016). Thus, maintaining the structure of the NL and NPCs is required for regulating a wide range of nuclear functions including transcription, DNA replication, DNA damage repair, force transition and the bidirectional flow of materials between the nuclear and cytoplasmic compartments.

The major structural determinants of the NL are the type-V intermediate filament proteins, the nuclear lamins (Goldman et al., 1986; McKeon et al., 1986). The lamins are classified as A-types (lamins A (LA) and C (LC)) and B-types (lamins B1 (LB1) and B2 (LB2)) (Fig. 1 A). LA and LC are derived from the *LMNA* gene by alternative splicing (Lin and Worman, 1993), whereas LB1 and LB2 are encoded by *LMNB1* and *LMNB2*, respectively (Biamonti et al., 1992; Höger et al., 1990; Lin and Worman, 1995; Maeno et al., 1995). In recent years, the detailed structure of the NL has been revealed by three-dimensional structured illumination microscopy (3D-SIM) combined with computer vision analysis and cryo-electron tomography (cryo-ET) (Shimi et al., 2015; Turgay et al., 2017). The lamin isoforms assemble into filamentous meshworks comprised of aggregates of filaments with a diameter of ∼3.5 nm in mouse embryonic fibroblasts (MEFs). These lamin filaments are non-randomly distributed into a layer with a mean thickness of ∼14 nm (Turgay et al., 2017). Notably, the four lamin isoform appears to assemble into distinct meshworks, each with a similar structural organization (Shimi et al., 2015). Knockdown (KD) and knockout (KO) of LB1 induce the formation of LA/C-rich structures on the nuclear surface including NE plaques and protrusions (Kittisopikul et al., 2021; Shimi et al., 2008).

**Figure 1.**
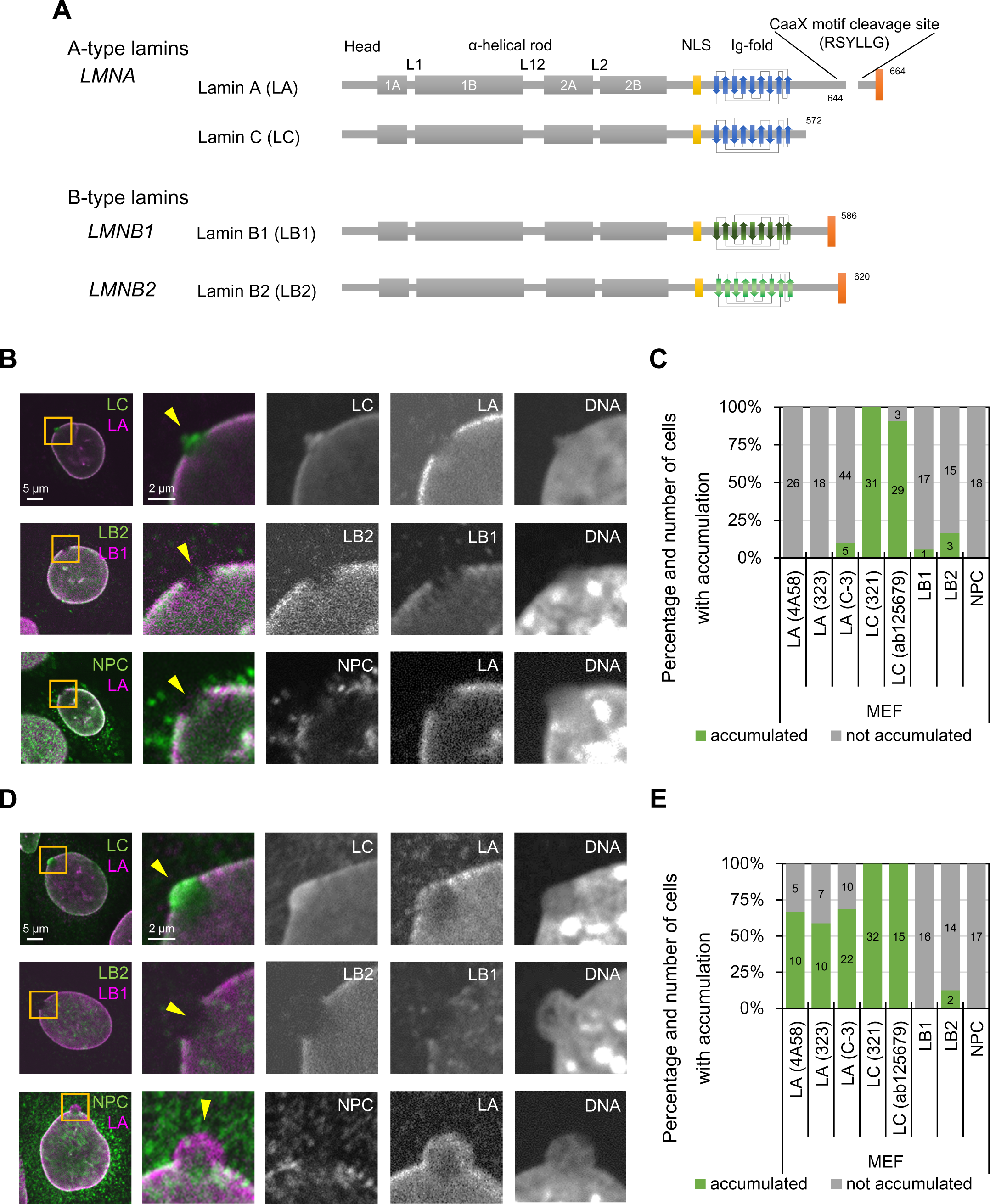
Difference of lamin isoforms in the structure and the accumulation kinetics at sites of NE rupture induced by laser microirradiation. **(A)** Protein architecture of lamin isoforms. The coiled-coil central rod domain (gray), the NLS (yellow), the β-strands comprising the Ig-fold (blue or green), and the CaaX motif box (red) are shown. **(B-E)** A 2-μm diameter spot at the NE in MEFs was laser-microirradiated to induce NE rupture, fixed within 10 min **(B and C)** or 60-70 min after laser microirradiation **(D and E)**, and then stained with a combination of anti-mouse and anti-rabbit antibodies, followed with Alexa Fluor 488-labeled anti-mouse or rabbit IgG and Cy5-labeled anti-rabbit or mouse IgG, and Hoechst 33342 for DNA. **(B and D)** Representative images of single confocal sections. Magnified views of the indicated areas by orange boxes are shown (the second to fifth columns). Color-merged images (the first and second columns) show anti-LC (321, green)/anti-LA (4A58, magenta), anti-LB2 (EPR9701(B), green)/anti-LB1 (B-10, magenta), and NPC (mAb414, green)/anti-LA (323, magenta). The ruptured sites are indicated with yellow arrowheads (the second columns). **(C and E)** Ratios of cells with (green) and without (gray) enrichments of the indicated antibodies at the rupture sites. The numbers of analyzed cells are indicated in the bar charts. Bars: 5 μm (the first column) and 2 μm (the second to fifth columns).

The leakage of a nuclear protein containing a nuclear localization signal (NLS) following NE rupture is observed both under normal physiological conditions and in pathological situations such as cancer cell migration through confined micro-environments during metastasis (Denais et al., 2016; Raab et al., 2016). The ruptures appear to be caused by a weakening of the structural integrity of the NE attributable to several factors including a loss of NE constituents (Chen et al., 2018), mechanical compression of cells (Hatch and Hetzer, 2016), tension applied directly to the NE (Zhang et al., 2019), and/or loss of certain tumor suppressors (Yang et al., 2017). At the same time, damaged regions of nuclear DNA in the vicinity of the rupture sites are sensed by the DNA sensors cyclic GMP-AMP synthase (cGAS) and its downstream signaling effector STING (Denais et al., 2016; Raab et al., 2016). The endosomal sorting complex required for transport III (ESCRT III) and Barrier-to-autointegration factor (BANF1/BAF) recruit LAP2-emerin-MAN1 (LEM) domain-containing INM proteins to the rupture sites (Denais et al., 2016; Halfmann et al., 2019; Raab et al., 2016). Subsequently, DNA repair factors accumulate at the DNA regions adjacent to the rupture sites to repair any DNA damage resulting from the rupture (Denais et al., 2016; Xia et al., 2019). The NE repair process is essential for the prevention of dysregulated nuclear functions due to DNA damage accumulation and the leakage of macromolecules to the cytoplasm (Denais et al., 2016; Raab et al., 2016).

The frequency of spontaneous NE rupture is significantly increased by the depletion of LA/C and/or LB1 (Chen et al., 2018; Chen et al., 2019; Earle et al., 2020; Kim et al., 2021; Robijns et al., 2016; Vargas et al., 2012). Numerous mutations have been found throughout the *LMNA* gene that cause a spectrum of human genetic disorders, collectively called laminopathies. It has been also reported that some of laminopathy mutations associated with dilated cardiomyopathy (DCM), muscular dystrophy (MD), familial partial lipodystrophy (FPLD), Limb-girdle muscular dystrophy type 1B (LGMD1B) and Hutchinson-Gilford Progeria syndrome (HGPS) frequently cause spontaneous NE rupture (De Vos et al., 2011; Earle et al., 2020; Kim et al., 2021). Thus, there is a large body of evidence supporting roles for the different lamin isoforms in protecting the NE from rupture under a wide range of physiological and pathological circumstances. However, it has yet to be determined if the lamins are actively involved in repair of NE ruptures. Here, we perform immunofluorescence for snapshot analyses and live-cell imaging for time-lapse analyses to determine the localization and dynamics of lamins after NE rupture in wild-type (WT), lamin-KO MEFs and *Lmna*-KO MEFs ectopically expressing mutant LC, containing known laminopathy mutations. Our data demonstrate that LA and LC have unique and important functions in repairing the ruptured NE.

## Results

### LA, LB1, LB2, and LC differentially respond to NE rupture

Despite previous studies by immunofluorescence to indicate that LA/C but not LB1 accumulated at sites of NE rupture (Denais et al., 2016; Xia et al., 2019), it is not known exactly which lamin isoforms are immediately targeted to the rupture sites. We therefore damaged a small region of the NE by 405-nm laser microirradiation to induce a rupture (Halfmann et al., 2019) in WT MEFs stably expressing super-folder Cherry harboring two NLSs derived from SV40 large T antigen and c-Myc (NLS-sfCherry). The cells were fixed within 10 min after laser microirradiation and stained with Hoechst 33342 for DNA and different combinations of specific antibodies directed against LA, LB1, LB2, LC, and NPCs. The site of NE rupture, often associated with a DNA protrusion containing decondensed chromatin, was enriched in LC and devoid of LA, LB1, LB2 and NPCs (Fig. 1 B, arrows; 1 C). The presence of LC but not LA at the rupture sites was consistently observed with different sources of the antibodies (Fig. 1 B, C and S1 A). However, when cells were fixed 60-70 min after laser microirradiation, both LA and LC were detected at the rupture sites (in ∼65% and 100% of nuclei for LA and LC, respectively; Fig. 1 D, E and Fig. S1 A). LB1, LB2, and NPCs remained absent from the rupture sites (Fig. 1 B-E). DNA protrusions were further pronounced in cells fixed 60-70 min after laser microirradiation (Fig. 1 B and D). Similar results were obtained using other cell lines stably expressing NLS-sfCherry, including those previously used for the demonstration of NE rupture (Earle et al., 2020; Halfmann et al., 2019), such as mouse myoblasts C2C12, hTERT-immortalized human foreskin fibroblasts BJ-5ta, and non-malignant breast epithelial cells MCF10A (Fig. S1 B-E).

### LC but not LA, LB1, or LB2 rapidly accumulates at the rupture sites

To follow the rapid repair process after NE rupture (Fig. 1 B-E and Fig. S1 A-E), we performed live-cell imaging of lamin isoforms, in accordance with a previous report (Halfmann et al., 2019). The mEmerald-fused LA, LB1, LB2, and LC were transiently expressed at low levels in WT MEFs expressing NLS-sfCherry and their localizations in response to NE rupture were analyzed by time-lapse confocal microscopy. After the induction of NE rupture by laser microirradiation, mEmerald-LA, LB1, and LB2 did not recover for at least ∼180 s (Fig. 2 A and B). In contrast, mEmerald-LC accumulated at the rupture sites within ∼50 s (Fig. 2A and B), where a NE plaque was formed before the protrusion from the nuclear main body.

**Figure 2.**
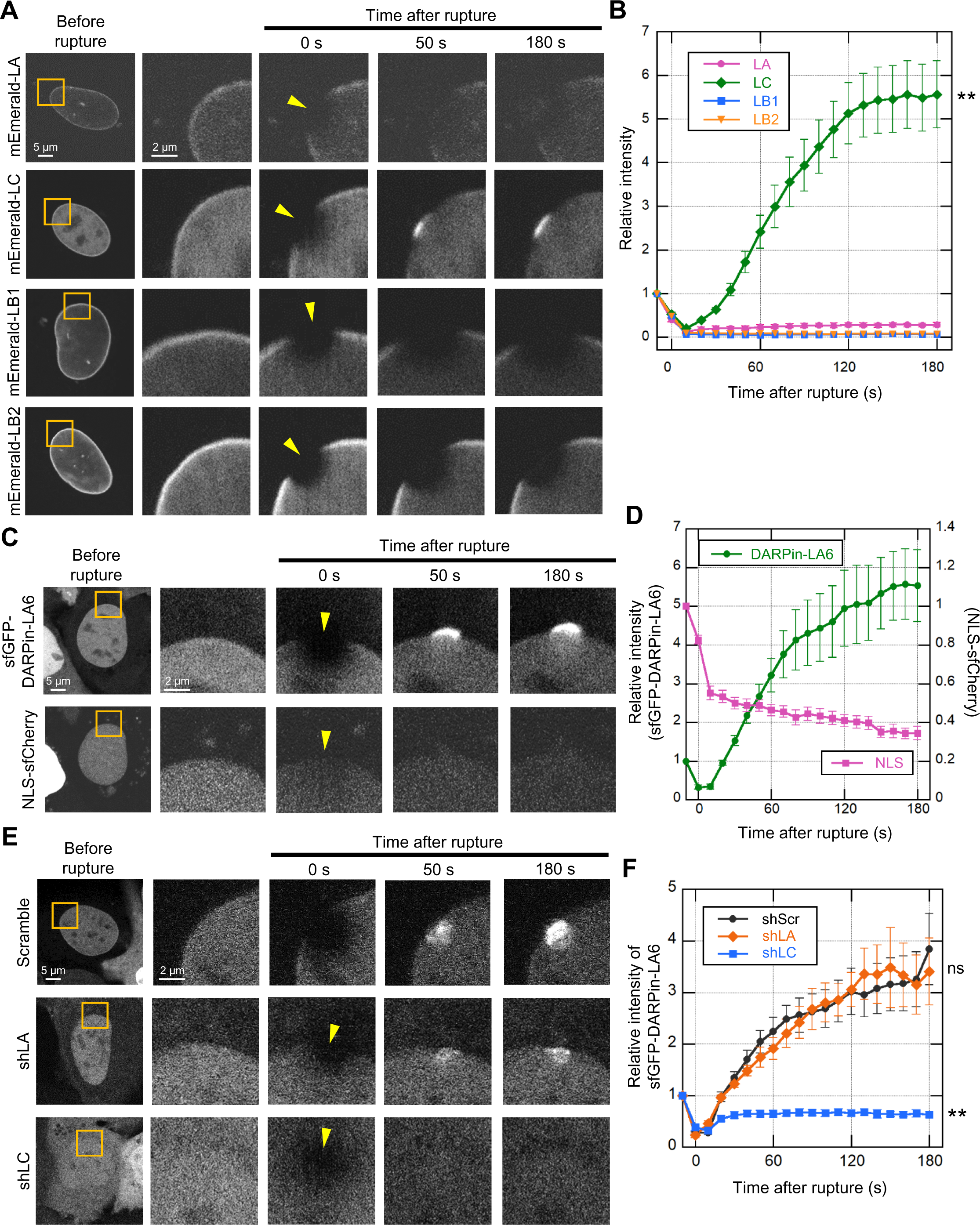
Rapid accumulation of mEmerald-LC at the rupture sites. During time-lapse imaging of exogenous LA, LC, LB1 and LB2 and endogenous LA and LC with 10 s intervals, a 2-μm diameter spot was laser-microirradiated to induce NE rupture (yellow arrowheads). **(A)** Dynamics of mEmerald-LA, LC, LB1 and LB2 in response to NE rupture in MEFs. **(B)** Fluorescence intensities of these mEmerald-lamins at the rupture sites were measured and normalized to the initial intensities. The graph represents means ± SEM (*n* = 20 cells from two independent experiments; **, P < 0.001 from others by a linear mixed model). **(C)** Dynamics of sfGFP-DARPin-LA6 and NLS-sfCherry in response to NE rupture in MEFs. **(D)** The sfGFP-DARPin-LA6 intensity at the rupture sites and NLS-sfCherry intensity in the nucleoplasm were measured and the relative intensities are plotted (means ± SEM; *n* = 20 cells from two independent experiments). **(E)** Dynamics of sfGFP-DARPin-LA6 in response to NE rupture in MEFs expressing shRNAs, scrambled control (shScr), shLA or shLC. **(F)** The sfGFP-DARPin-LA6 intensity at the rupture sites was measured and the relative intensities are plotted (means ± SEM; *n* = 10 cells; **, P < 0.001; ns, P > 0.05 from control by a linear mixed model). Bars: 5 μm (the first column) and 2 μm (the second to fifth columns).

To confirm that the dynamics of endogenous LA and LC are represented by the ectopic expression of mEmerald-LA and -LC, these isoforms were directly labeled with a LA/C-specific genetically encoded probe, DARPin (designed ankyrin repeat protein) (Zwerger et al., 2015), fused with super-folder GFP (sfGFP). To avoid interfering with lamin assembly and functions, among all the DARPins we chose LaA_6, which binds moderately to the head domain of LA/C (Kd = 8.25×10^−7^ M) (Zwerger et al., 2015), to construct sfGFP-DARPin-LA6. In WT MEFs expressing NLS-sfCherry, the sfGFP-DARPin-LA6 signals were detected throughout the nucleus probably due to its low affinity for LA/C (Fig. 2 C). After laser microirradiation, sfGFP-DARPin-LA6 accumulated at the rupture sites within ∼50 s and its accumulation increased for at least 180 s (Fig. 2 C and D). At the same time, the fluorescence intensity of NLS-sfCherry in the nucleus was decreased due to leakage into the cytoplasm (Fig. 2 C and D).

Next, to examine which lamin isoform, LA or LC, contributes to sfGFP-DARPin-LA6 accumulation at the rupture sites, we employed LA- and LC-specific KD by lentivirus-mediated expression of short hairpin RNAs (shRNAs) that selectively target LA or LC (shLA or shLC) (Harr et al., 2015; Wong et al., 2021). The expression of shLA and shLC successfully reduced their target isoforms (Fig. S2 A and B). In cells expressing scrambled control or shLA, sfGFP-DARPin-LA6 accumulated at the rupture sites (Fig. 2 E and F), as observed in non-treated cells (Fig. 2 C and D). In contrast, the accumulation of sfGFP-DARPin-LA6 was significantly reduced with shLC (Fig. 2 E and F). These data are consistent with the observations by immunofluorescence and mEmerald-lamins, supporting the view that endogenous LC but not LA rapidly accumulates at the rupture sites.

LA/C are known to contribute to the prevention of NE rupture under mechanical stress (Denais et al., 2016; Halfmann et al., 2019; Raab et al., 2016). Because LC accumulated at the rupture sites in a LA-independent manner (Fig. 2 E and F), LC could slow the leakage of nuclear proteins without LA. To test this idea, HaloTag harboring NLS derived from SV40 large T antigen (NLS-Halo) was expressed in scrambled control and LC-KD MEFs, and its nuclear leakage kinetics in responses to NE rupture was analyzed by time-lapse confocal microscopy. The cytoplasmic-to-nuclear intensity (C/N) ratio of NLS-Halo after the rupture was rapidly increased in LC-KD cells compared to the control cells (Fig. S2 C and D). This suggests that LC alone can function in slowing the leakage from the rupture site.

### LC is recruited from the nucleoplasm to the rupture sites

LC harbors a unique six amino acid segment at the C-terminus following the amino acid sequence shared by LA and LC. To examine if the LC-specific segment is required for the rapid accumulation at the rupture sites, a deletion mutant of LC fused with mEmerald that lacks the six amino acids (Δ567-572, or Δ6aa) was expressed in *Lmna*-KO MEFs expressing NLS-sfCherry and its accumulation at rupture sites was analyzed by live-cell imaging. We used the LA/C-null background for our mutant analysis to avoid complications by interactions between the mutant and endogenous LC proteins. The Δ6aa mutant accumulated at the rupture sites, similarly to the full-length LC (Fig. S3 A-C), indicating that the different dynamics of LA and LC are not attributable to the LC-specific six amino acids.

Nucleoplasmic LA and LC, previously described as a part of a ‘nucleoplasmic veil’ (Moir et al., 2000), are highly diffusible in the nucleoplasm compared to those at the NL (Broers et al., 1999; Shimi et al., 2008). Because LC is more abundant in the nucleoplasm and more detergent-extractable than LA (Kolb et al., 2011; Markiewicz et al., 2002; Wong et al., 2021), the nucleoplasmic pool of LC could readily diffuse to the rupture sites. To test this idea, mEmerald-LC was photobleached in the nucleoplasm of *Lmna*-KO MEFs expressing NLS-sfCherry, and then NE rupture was immediately induced by laser microirradiation. The mEmerald-LC signals were exclusively observed in the NL after photobleaching, and the rupture sites remained devoid of mEmerald-LC for at least 180 s (Fig 3 A and B), indicating that LC accumulated at the rupture sites originated from the nucleoplasmic pool and not from the NL. To examine if the different accumulation kinetics of LA and LC are attributable to the different levels of nucleoplasmic pools of LA and LC, we used cells that overexpress LA while we routinely used those with low expression (Fig. 2 A and B). In the highly expressing cells, mEmerald-LA modestly accumulated at the rupture sites (∼1.2-fold enrichment 150 s after irradiation; Fig. S3 D-F). Thus, the diffusible LA and LC in the nucleoplasm appear to be involved in the accumulation.

**Figure 3.**
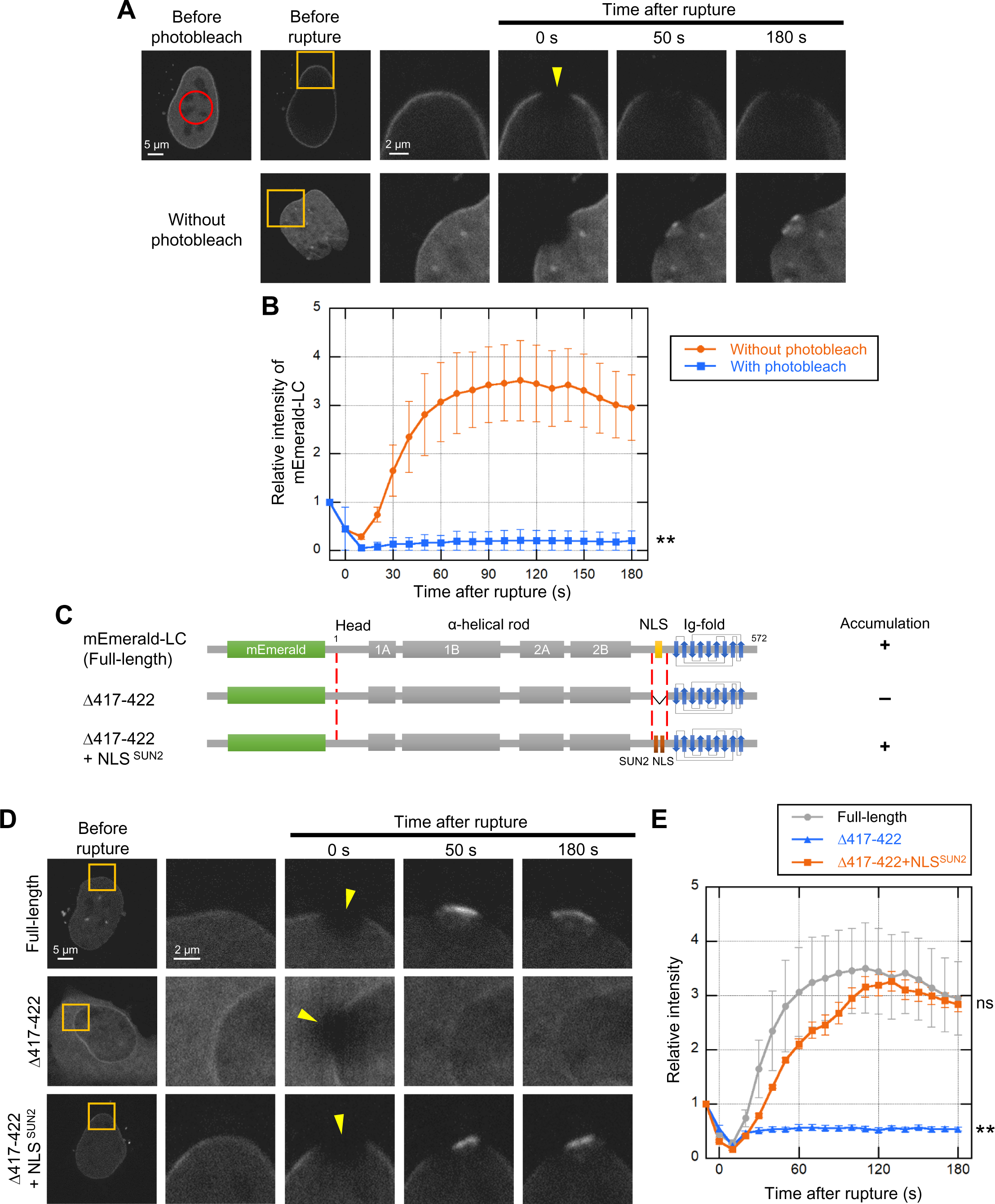
Rapid accumulation of nucleoplasmic LC at the rupture sites. **(A)** Dynamics of mEmerald-LC in response to NE rupture with or without photobleaching the nucleoplasmic fraction. The right four columns are magnified views of orange boxes. (Top row) A nucleoplasmic area in *Lmna*-KO MEFs expressing mEmerald-LC (red circle) was photobleached using 488-nm laser, and then a 2-μm spot at the NE (yellow arrowhead) was microirradiated using 405-nm laser during time-lapse imaging with 10 s intervals. (Bottom row) The control cells without photobleaching. Bars: 5 μm (the left two columns) and 2 μm (the right four columns). **(B)** Relative fluorescence intensity of mEmerald-LC at the rupture sites. The mEmerald-LC intensities relative to the initial point are plotted (means ± SEM; *n* = 20 cells from two independent experiments; **, P < 0.001 from without photobleaching by a linear mixed model). **(C**-**E)** Requirements of an NLS for LC accumulation at the rupture sites. mEmerald-LC full-length, Δ417-422 (ΔNLS) and Δ417-422 + NLS^SUN2^ (ΔNLS+sunNLS) were expressed in *Lmna*-KO MEFs and the NE rupture assay was performed as in **A** and **B**, without pre-photobleaching. **(C)** Architecture of the mEmerald-LC NLS mutants. The summary of their dynamics is indicated on the right (+, accumulated at the rupture site; -, not accumulated). **(D)** Dynamics of mEmerald-LC NLS mutants in response to NE rupture. Bars: 5 μm (the first column) and 2 μm (the second to fifth columns). **(E)** Relative fluorescence intensities of the mEmerald-LC NLS mutants (means ± SEM; *n* = 10 cells; **, P < 0.001; ns, P > 0.05 from full-length by a linear mixed model). Full-length (gray) is a reproduction of “Without photobleach” in **B**.

The transport of the lamins from the cytoplasm to the nucleus is mediated through their NLSs (Loewinger and McKeon, 1988). To test if the NLS of LC determines the abundance of the nucleoplasmic pool for the rapid accumulation at the rupture sites, NLS mutants of LC fused with mEmerald were expressed in *Lmna*-KO MEFs expressing NLS-sfCherry (Fig. 3 C). An NLS-deletion mutant (Δ417-422; ΔNLS), which mostly remained in the cytoplasm compared to the full-length LC, did not accumulate at the rupture sites (Fig. 3 D and E), suggesting that the nuclear localization of LC is critical for the accumulation. However, the LC NLS sequence, rather than the nuclear localization, could have a specific function in the accumulation. To test this possibility, we expressed another NLS mutant of LC, in which the NLS is replaced with an NLS from a component of the LINC complex, SUN2 (Turgay et al., 2010) (Δ417-422 + NLS^SUN2^; ΔNLS+sunNLS). This mutant localized to the NL and the nucleoplasm, and accumulated at the rupture sites (Fig. 3 D and E), indicating that the abundance of the nucleoplasmic pool of LC, but not the NLS per se, is critical for the accumulation.

### Accumulation of LC at the rupture sites requires the Ig-fold domain

BAF binds to the LEM domain of INM proteins (Lee et al., 2001) and recruits them to rupture sites (Halfmann et al., 2019). Because BAF also binds to the immunoglobulin (Ig)-like fold (Ig-fold) domain in the C-terminal tail of LA/C but not LB1 (Samson et al., 2018), the Ig-fold might mediate the recruitment of LC to the rupture sites through an interaction with BAF. We therefore tested this idea by expressing Ig-fold mutants of LC fused with mEmerald in *Lmna*-KO MEFs expressing NLS-sfCherry (Fig. 4 A). LC lacking the Ig-fold (Δ432-572, or ΔTail), which corresponds to the Q432X mutation found in DCM patients (Møller et al., 2009), failed to accumulate at the rupture sites (Fig. 4 B and C). Replacing the Ig-fold with that of LB1 (Δ432-548 + Ig-fold^LB1^; ΔIgF+b1IgF) also did not accumulate (Fig. 4 B and C).

**Figure 4.**
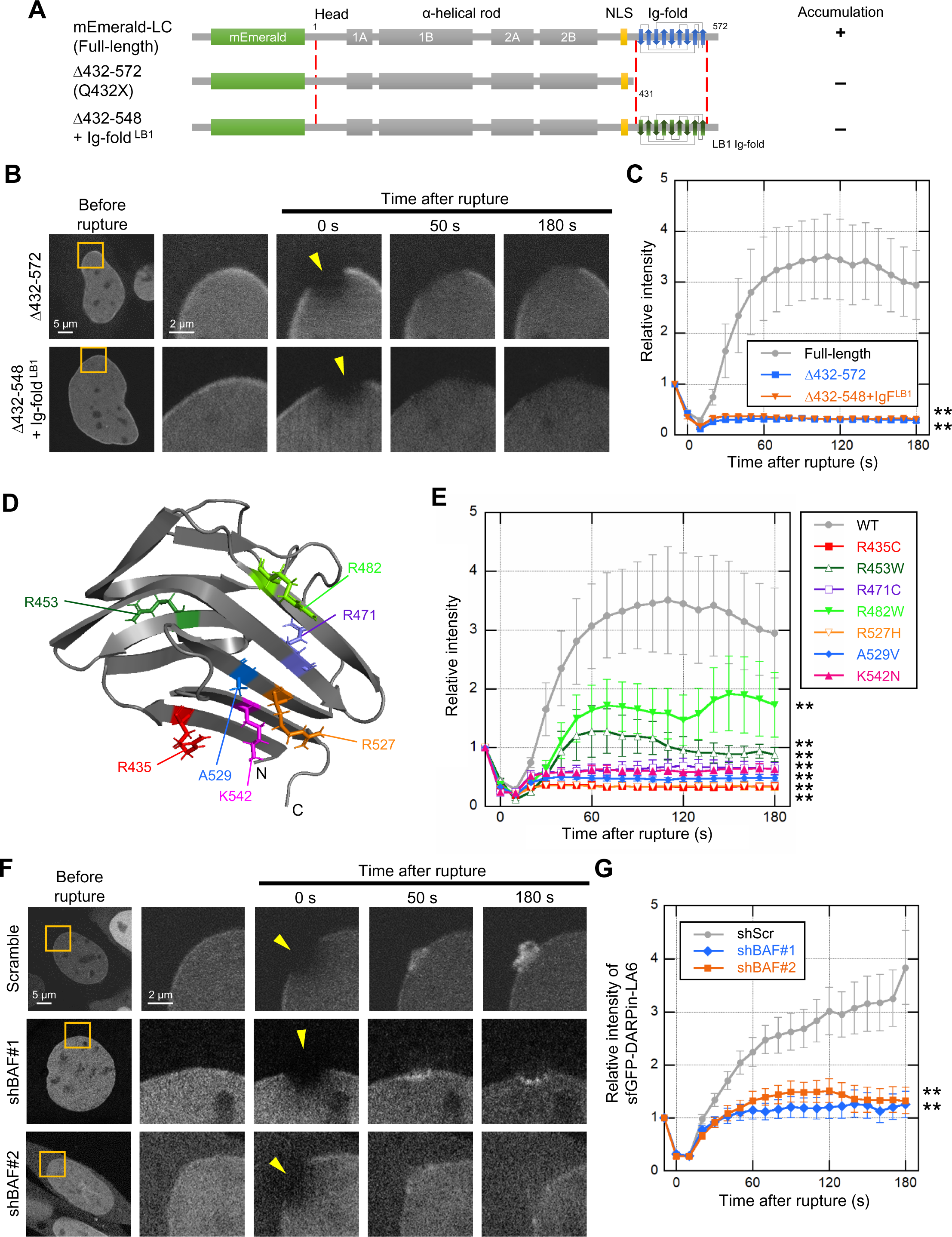
Effect of LC Ig-fold laminopathy mutations and BAF KD on accumulation kinetics of LC at the rupture sites. The NE rupture assay was performed with mEmerald-LC mutants in *Lmna*-KO MEFs (**A**-**E**) and sfGFP-DARPin-LA6 in BAF-KD MEFs (**F** and **G**). **(A)** Architecture of mEmerald-LC full-length, Δ432-572 (ΔTail) and Δ432-548 + Ig-fold^LB1^ (ΔIgF+b1IgF). The summary of their dynamics is indicated on the right (+, accumulated at the rupture site; -, not accumulated). **(B)** Dynamics of mEmerald-LC Ig-fold mutants in response to NE rupture in *Lmna*-KO MEFs. The right four columns are magnified views of orange boxes. **(C)** Relative fluorescence intensities of the mEmerald-LC Ig-fold mutants (means ± SEM; *n* = 10 cells; **, P < 0.001 from full-length by a linear mixed model). **(D)** Positions of laminopathy mutations in the LA/C Ig-fold structure (PDB ID 1IFR). The amino acid residues whose mutations affect BAF binding affinity in vitro (Samson et al., 2018) are colored (red, no detectable binding; orange and magenta, very weak binding; and purple and blue, ∼5-fold weaker binding to the WT). The two residues whose mutations have no effect on BAF binding affinity *in vitro* are shown in dark green and light green. **(E)** Relative fluorescence intensities of the mEmerald-LC Ig-fold laminopathy mutants in *Lmna*-KO MEFs (means ± SEM; *n* = 10 cells; **, P < 0.001 from full-length by a linear mixed model). See Fig. S4 A for microscopic images. **(F)** Dynamics of sfGFP-DARPin-LA6 in response to NE rupture in WT MEFs expressing shRNAs, scrambled control (shScr), shBAF#1 or shBAF#2. The right four columns are magnified views of orange boxes (See Fig. S4 B and C for the validation of KD by immunofluorescence and immunoblotting). **(G)** Relative fluorescence intensities of sfGFP-DARPin-LA6 in the indicated cells (means ± SEM; *n* = 10 cells; **, P < 0.001 from the shScramble by a linear mixed model). (**C** and **E**) Full-length (gray) is a reproduction of “Without photobleach” in Fig. 3 B. (G) shScr (gray) is a reproduction of “shScramble” in Fig. 2 F. (**B** and **F**) Bars: 5 μm (the first column) and 2 μm (the second column to others).

Some of laminopathy mutations (R435C, R453W, R471C, R482W, R527H, A529V, and K542N) that reside within the Ig-fold domain cause cardiac and skeletal muscle diseases, dysplasia, and progeroid syndrome (Fig 4 D, and Table 1). When expressed in *Lmna*-KO MEFs expressing NLS-sfCherry, all mutant fused with mEmerald were localized to the NL and the nucleoplasm (Fig. S4 A). Five of these mutants (R435C, R471C, R527H, A529V, and K542N) that show low binding affinity to BAF in vitro (Table 1) (Samson et al., 2018) did not accumulate at the rupture sites for at least 180 s (Fig. 4 E). The other two mutants (R453W and R482W), which bind to BAF in vitro with similar affinities to the WT (Table 1), slowly accumulated at the rupture sites compared to WT LC (Fig. 4 E). This data suggests that the LC binding with BAF mediated through the Ig-fold is important in the accumulation.

**Table 1.**
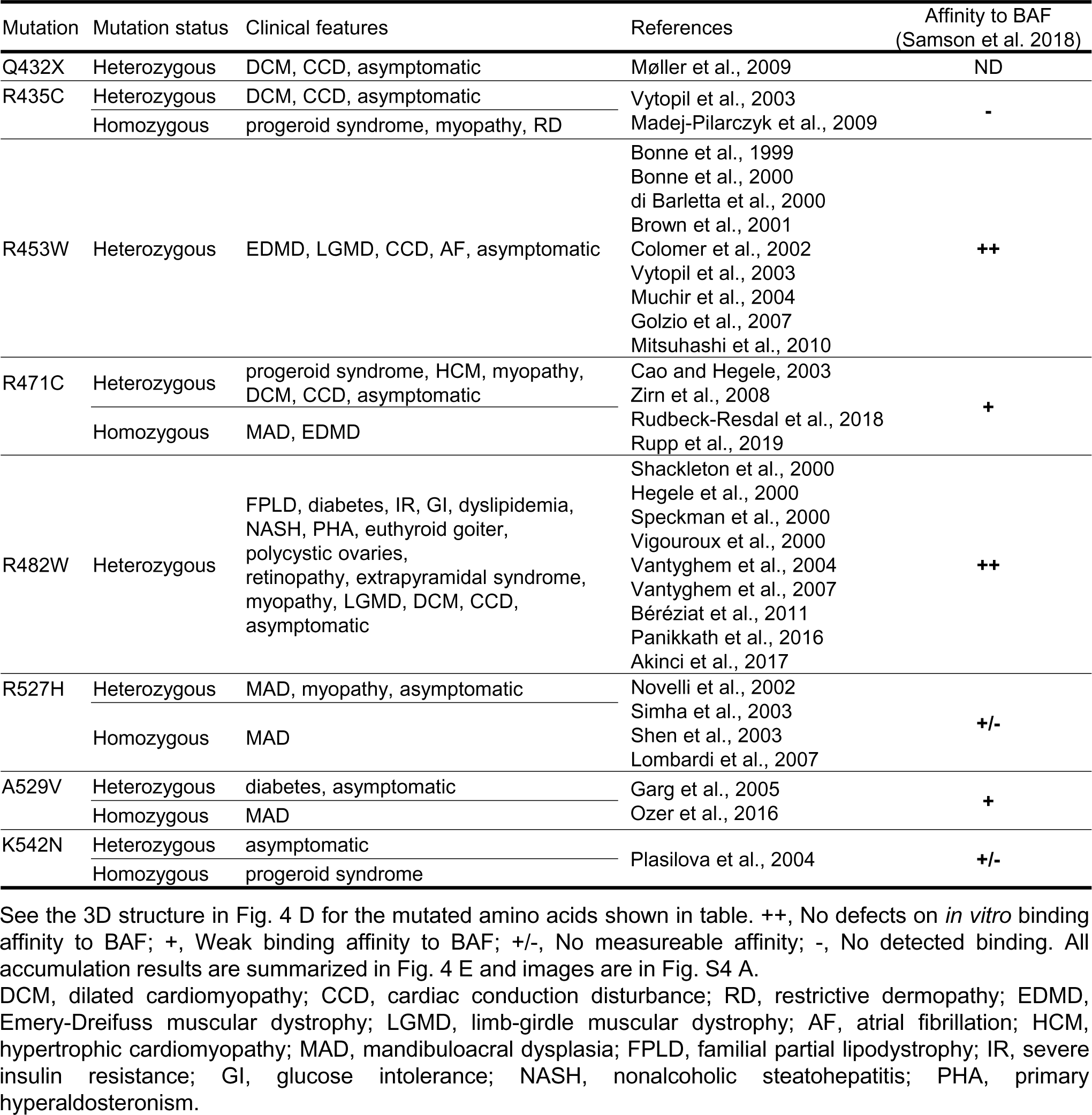
Clinical features of Laminopathy mutations on tail region of LMNA gene

Next, the role of BAF in recruiting LC to the rupture sites was examined by BAF-KD in WT MEFs. The accumulation of sfGFP-DARPin-LA6 at the rupture sites was significantly diminished in BAF-KD cells (Fig. 5 F and G). While sfGFP-DARPin-LA6 was observed in the DNA protruded regions in control cells (Fig. 5 F, top row), it was only localized to the peripheries of the nuclear main bodies in BAF-KD cells (Fig. 5F, the second row). To confirm this observation, we fixed cells within 3 min after laser microirradiation and stained DNA with Hoechst 33342. sfGFP-DARPin-LA6 signals were indeed detected in protruded DNA in control cells but not in BAF-KD cells (Fig. S4 D). From these data, it can be concluded that BAF is required for LC accumulation at the protruded DNA regions.

**Figure 5.**
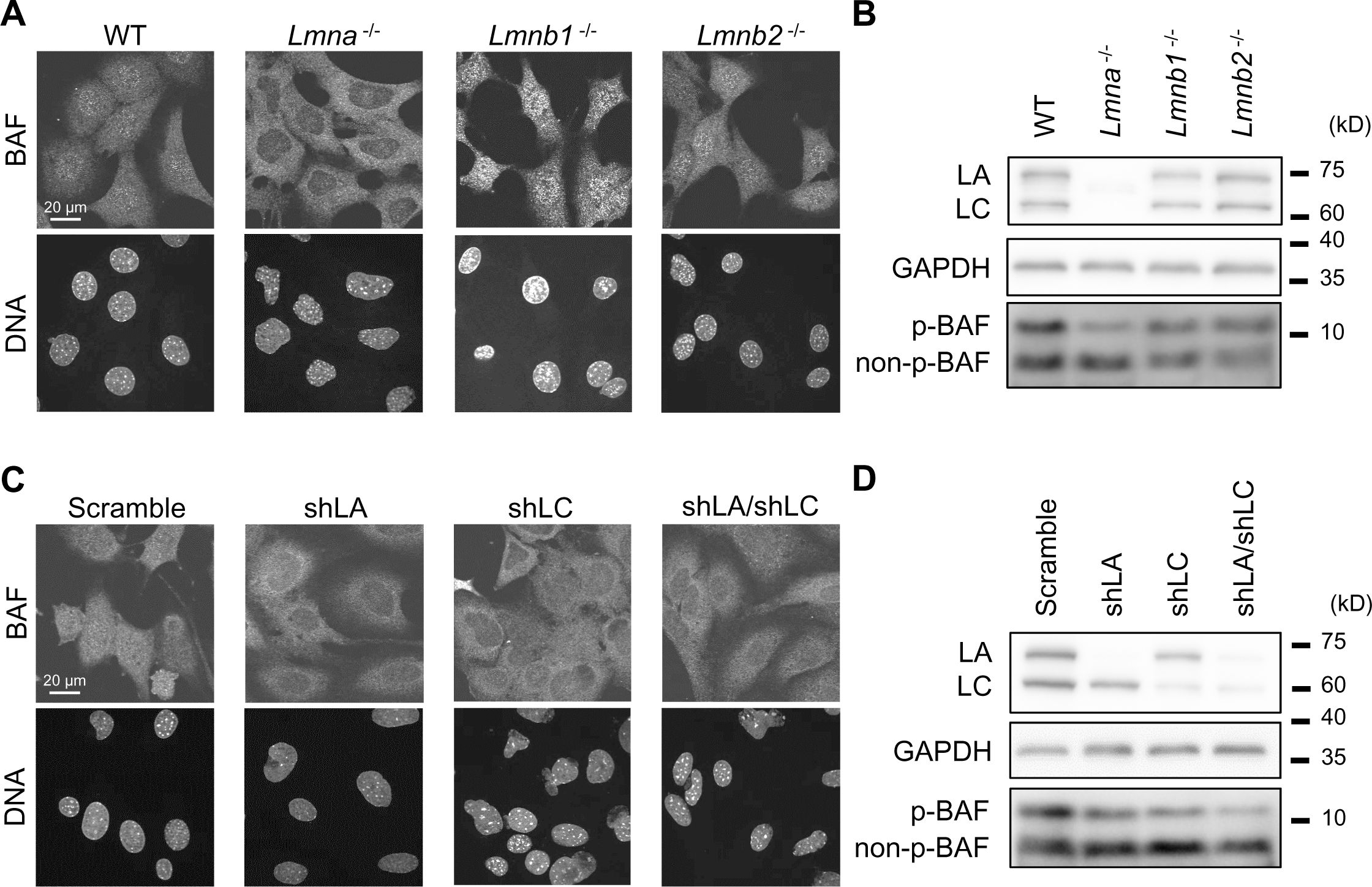
Effect of lamin depletion on localization and phosphorylation of BAF. **(A and B)** The localization and phosphorylation of BAF in WT, *Lmna* ^-/-^, *Lmnb1* ^-/-^, and *Lmnb2* ^-/-^ MEFs was analyzed by immunofluorescence (**A**) and immunoblotting, respectively (**B**). **(A)** Single confocal sections of the indicated cells stained with anti-BANF1/BAF (EPR7668), followed with Alexa Fluor 488-labeled anti-rabbit IgG, and Hoechst 33342 for DNA. **(B)** Whole cell lysates from the indicated cells were probed with anti-LA/C, anti-GAPDH (as loading control), and anti-BANF1/BAF (EPR7668). **(C** and **D)** The localization and phosphorylation of BAF in WT MEFs expressing scrambled control, shLA, shLC or a combination of shLA and shLC was analyzed by immunofluorescence (**C**) and immunoblotting, respectively (**D**). **(C)** Single confocal sections of the indicated cells stained with anti-BANF1/BAF (EPR7668), followed with Cy5-labeled anti-rabbit IgG, and Hoechst 33342 for DNA. **(D)** Whole cell lysates from the indicated cells were probed with anti-LA/C, anti-GAPDH (as loading control), and anti-BANF1/BAF (EPR7668). (**A** and **C**) Bars: 20 μm. (**B** and **D**) Positions of the size standards are shown on the right.

### LA and LC facilitate BAF localization to the nucleus

It has been reported that LA and/or LC play a role in retaining BAF inside the nucleus (Lin et al., 2020). Therefore, we examined if BAF localization is affected by depletion of specific lamins by staining endogenous BAF in WT, *Lmna*-KO, *Lmnb1*-KO and *Lmnb2*-KO MEFs using two different antibodies. The nuclear BAF signals were significantly decreased by *Lmna*-but not *Lmnb1*- or *Lmnb2*-KO (Fig. 5 A, and Fig. S5 A and B). Because cytoplasmic BAF is known to be less phosphorylated than nuclear BAF (Zhuang et al., 2014), *Lmna* KO could reduce the level of phosphorylated BAF (p-BAF), which can be identified as a retarded band by immunoblotting (Nichols et al., 2006). The p-BAF band intensity was decreased to ∼50% in *Lmna*-KO cells compared to WT cells without an increase in level of the non-phosphorylated form (non-p-BAF) whereas the reduction was marginal in *Lmnb1*- and *Lmnb2*-KO cells (Fig. 5 B).

To determine which lamin isoform, LA or LC, is responsible for the nuclear localization and phosphorylation of BAF, they were individually or simultaneously knocked down in WT MEFs using LA- and LC-specific shRNAs, and the combination of both. The nuclear BAF signals were significantly reduced by LA-, LC-, and LA/C-KD (Fig 5 C). The p-BAF band intensities were also decreased to ∼50%, ∼40% and ∼30% in LA-, LC-, and LA/C-KD cells, respectively, compared to the control cells (Fig. 5 D). These data indicate that both LA and LC are involved in the nuclear localization and phosphorylation of BAF.

The fact that the nuclear p-BAF level is reduced in *Lmna*-KO MEFs raised a possibility that the effect of LC-mutant expression is affected by such background. To test the effect of BAF levels, HaloTag-fused BAF (Halo-BAF) was co-expressed with mEmerald-fused LC (full-length) and the deletion mutants (ΔNLS and ΔTail) that did not accumulate at the rupture sites in *Lmna*-KO MEFs. The full-length LC accumulated at the rupture sites but the ΔNLS and ΔTail did not (Fig. S5 C and D), as observed before without BAF overexpression (Fig. 3 and 4). Thus, endogenous BAF is sufficient for the recruitment of ectopically expressed LC to the rupture sites in the LA/C-null background and an excess BAF does not rescue the defects in the mutants.

### Cytoplasmic cGAS accumulates at the rupture sites and affects LC and BAF accumulation

The DNA sensor cGAS can detect nuclear DNA at the ruptured sites (Denais et al., 2016; Halfmann et al., 2019; Raab et al., 2016). To examine a potential function of cGAS in the accumulation of LC and BAF at the rupture sites, sfCherry-fused cGAS (cGAS-sfCherry) was co-expressed with sfGFP-DARPin-LA6 and Halo-BAF in WT MEFs. cGAS-sfCherry exhibited either the preferential nuclear or cytoplasmic localization in different cells (Fig. 6 A and B). Immediately after laser microirradiation, sfGFP-DARPin-LA6 and Halo-BAF simultaneously accumulated at the ruptured sites, and with a short delay, cGAS-sfCherry entered the nucleus from the cytoplasm through the opening of the ruptured NE to accumulate (Fig. 6 A, C, and Fig. S5 C). In contrast, nuclear cGAS-sfCherry did not show any accumulation at all (Fig. 6 B, C, and Fig. S5 D). Similar results were obtained by immunofluorescence using cells fixed within 10 min after laser microirradiation. cGAS accumulation was significantly weak in cells with nuclear cGAS compared to cells with cytoplasmic cGAS (Fig. 6 D and E). These results are consistent with previous findings to indicate that cGAS is active in the cytoplasm whereas nuclear cGAS is inactive (Kujirai et al., 2020; Michalski et al., 2020; Zhao et al., 2020). The accumulation levels of sfGFP-DARPin-LA6 and Halo-BAF at the rupture sites were lower in cells with nuclear cGAS-sfCherry compared to those with the cytoplasmic cGAS-sfCherry (Fig. 6 C). DNA protrusion became obvious in cells with cytoplasmic cGAS-sfCherry compared to those with nuclear cGAS-sfCherry (Fig. 6 A and B), suggesting that cGAS accumulation at the rupture sites could facilitate DNA protrusion and induce more accumulation of LC and BAF.

**Figure 6.**
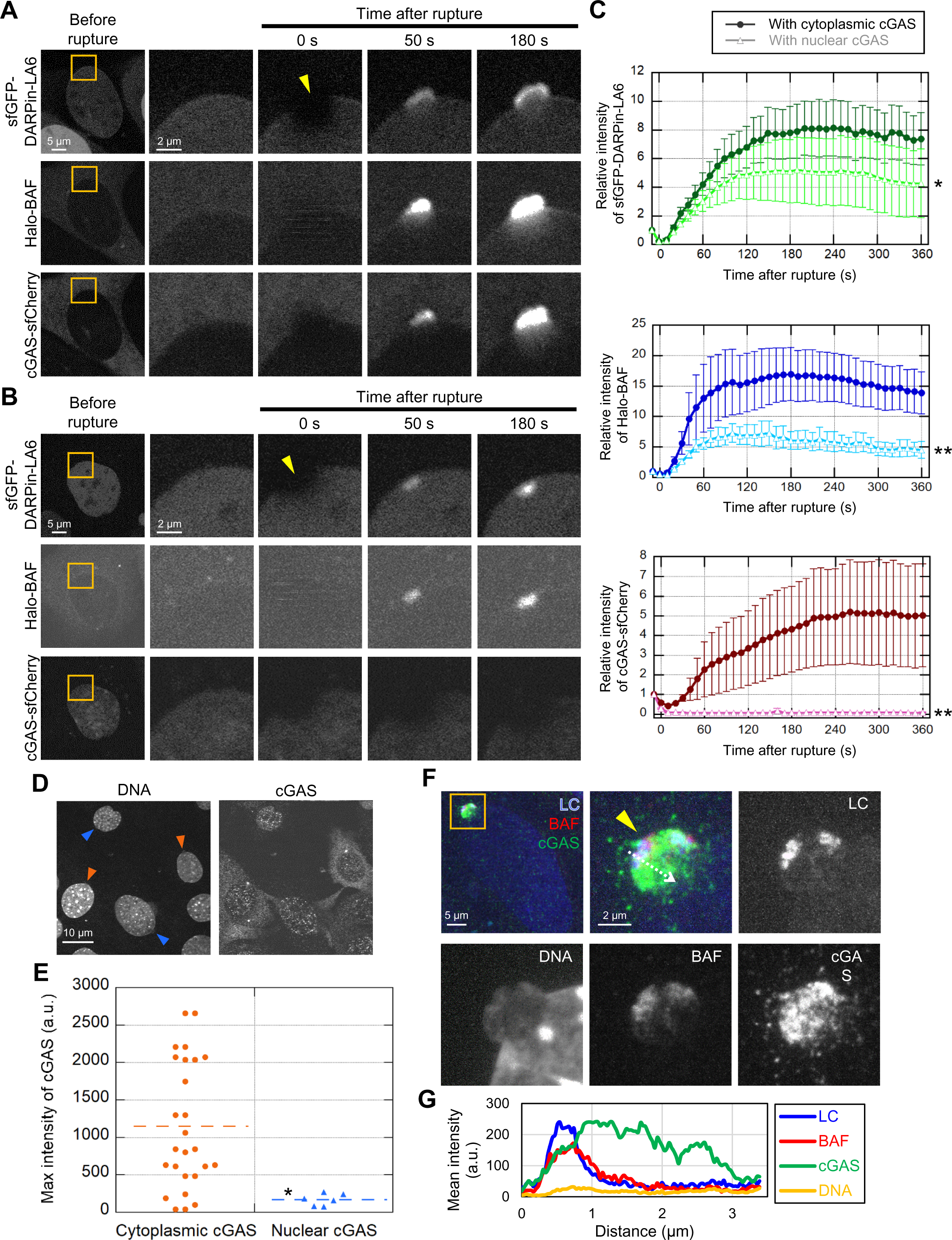
Dynamics of LC, BAF, and cGAS in response to laser microirradiation-induced NE rupture. **(A and B)** Dynamics of sfGFP-DARPin-LA6, Halo-BAF, and cGAS-sfCherry in response to NE rupture. cGAS-sfCherry was localized to the cytoplasm in some cells (**A**) or to the nucleus in the others (**B**). The right four columns are magnified views of orange boxes, and the rupture sites are indicated with yellow arrowheads. **(C)** Relative intensities of sfGFP-DARPin-LA6 (top), Halo-BAF (middle), and cGAS-sfCherry (bottom) at the rupture sites in cells with cytoplasmic or nuclear cGAS (means ± SEM; *n* = 10 cells; *, P < 0.01; **, P < 0.001 by a linear mixed model). **(D)** Representative images of single confocal sections of MEFs fixed within 10 min after laser microirradiation and stained with anti-cGAS, Alexa Fluor 488-labeled anti-rabbit IgG, and Hoechst 33342 for DNA. Colored arrowheads indicate sites of NE rupture induced by laser microirradiation in cells with cytoplasmic cGAS (orange) or nuclear cGAS (blue). Bar: 10 μm. **(E)** Fluorescence intensities of the cGAS accumulated at the rupture sites was measured. (*, P < 0.05 from cells with the accumulation of cytoplasmic cGAS by a Mann-Whitney U test). **(F)** Maximum intensity Z-projection of high-resolution confocal images of LC, BAF and cGAS in a NE protrusion from the nuclear main body. The sfGFP-DARPin-LA6 expressing cells are fixed within 10 min after laser microirradiation, stained with anti-BANF1 (3F10-4G12) and anti-cGAS, followed with Cy5-labeled anti-mouse IgG and Alexa Fluor 568-labeled anti-rabbit IgG, respectively, and Hoechst 33342 for DNA. Magnified views of an orange boxed area (top left) are shown as merged and individual images. Merged images (top left and top middle) show LC (blue), BAF (red), and cGAS (green). **(G)** Line intensity profiles over the NE protrusion. Fluorescence intensity on the white dotted-line arrow was measured and plotted. (**A**, **B** and **F**) Bars: 5 μm (**A** and **B**, the first column; and **F**, the top left); and 2 μm for the magnified views.

To visualize the localization of cGAS, BAF, and LA/C at a higher resolution, we stained endogenous BAF and cGAS in WT MEFs expressing sfGFP-DARPin-LA6 in cells that were fixed within 10 min after laser microirradiation. cGAS was often localized inside a protruded DNA region whereas LA/C and BAF were localized to the peripheries of protruded DNA regions (Fig. 6 F and G).

### LA/C is involved in the localization of BAF and cGAS after NE rupture

As both A- and B-type lamins are involved in protecting the NE from rupture (Denais et al., 2016; Halfmann et al., 2019; Raab et al., 2016; Young et al., 2020), we analyzed the dependence of BAF and cGAS accumulation at rupture sites on different lamin isoforms by immunofluorescence using WT MEFs, *Lmna*-, *Lmnb1*- and *Lmnb2*-KO MEFs expressing NLS-sfCherry. The expression levels and localization of cGAS were similar in all the MEFs (Fig. S5 G and H). In cells fixed within 10 min after laser microirradiation, the accumulation of BAF and cGAS was significantly reduced in *Lmna*-KO cells compared to WT and *Lmnb1*- and *Lmnb2*-KO cells (Fig. 7 A-C). Unlike DNA protrusions with BAF and cGAS in WT cell nuclei, laser-microirradiated DNA was located in the nuclear interior of the *Lmna*-KO cell nuclei, as indicated with the weak signals (Fig. S5 I). In *Lmnb1*-KO cells, BAF- and cGAS-positive protrusions were often observed without laser microirradiation-induced rupture (Fig. 7 A), as previously reported (Vergnes et al., 2004; Young et al., 2020). Taken together with the data above, these results suggest that LA/C, BAF, and cGAS are concertedly accumulated at the NE rupture sites.

**Figure 7.**
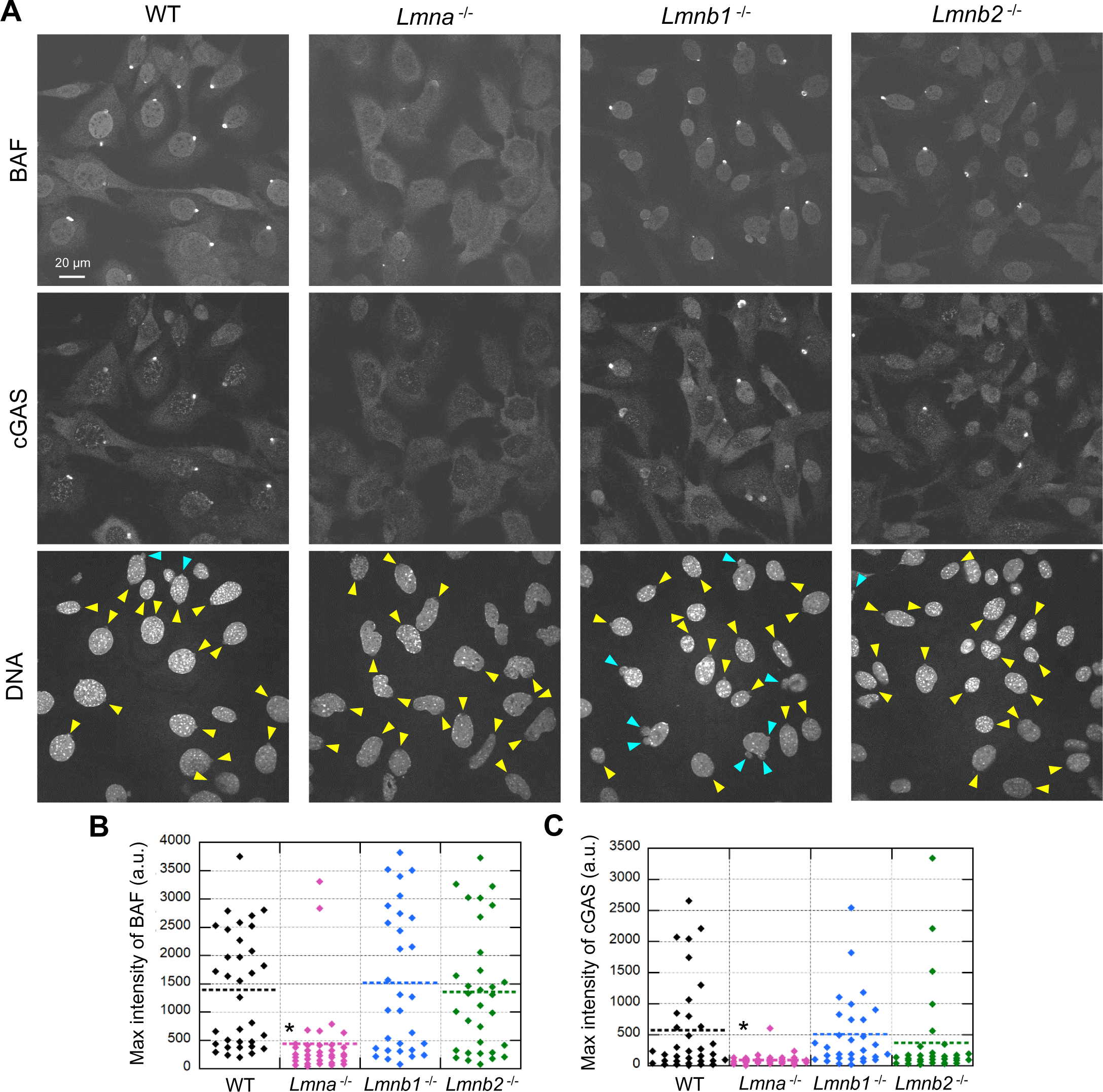
Effect of lamin depletion on BAF and cGAS accumulation at the rupture sites. **(A)** Single confocal sections of WT, *Lmna* ^-/-^, *Lmnb1* ^-/-^, and *Lmnb2* ^-/-^ MEFs fixed within 10 min after laser microirradiation and stained with anti-BANF1 (3F10-4G12) and anti-cGAS, followed with Alexa Fluor 488-labeled anti-rabbit IgG, Cy5-labeled anti-mouse IgG, and Hoechst 33342 for DNA. Yellow arrowheads indicate laser microirradiation-induced NE rupture sites. Blue arrowheads indicate spontaneously produced NE protrusions. Bar: 20 μm. (B and C) The max intensities of BAF (B) and cGAS (C) signals at the rupture sites. The plotted points are from two independent experiments (*n* = 36, 35, 32, and 34 for WT, *Lmna* ^-/-^, *Lmnb1* ^-/-^, and *Lmnb2* ^-/-^, respectively; *, P < 0.05 from others by a Steel-Dwass multiple comparison).

## Discussion

When the NE is locally ruptured under various circumstances, NE components are recruited to the rupture sites with the ESCRT III complex and BAF for repair, and the DNA regions adjacent to the rupture sites are sensed with cGAS (Denais et al., 2016; Halfmann et al., 2019; Raab et al., 2016). Lamins can protect the NE from rupture (Chen et al., 2018), but the specific role of lamins in repairing the ruptured NE and sensing the adjacent DNA regions remain largely unknown. In this study, we have analyzed the dynamics of lamin isoforms, BAF, and cGAS in the early response to NE rupture. Fig. 8 summarizes our findings and those of previous studies. The diagrams depict three time points; before the NE rupture and ∼1 min and ∼10 min after laser microirradiation. Under normal conditions without rupture, LA and LC form complexes with p-BAF in the NL and the nucleoplasm. LC is more abundantly present in the nucleoplasm than LA. The non-p-BAF and cGAS are present in the cytoplasm. Upon the induction of NE rupture (∼1 min, middle), diffusible LC-p-BAF complexes rapidly accumulate at the rupture sites to form a NE plaque. The non-p-BAF and cGAS can access protruded DNA regions because of the opening of the ruptured NE. At a later time (∼10 min, bottom), DNA protrusion becomes more evident with the further accumulation of both LA and LC, p-BAF, and cGAS. B-type lamins are likely to return to the rupture sites at a later time point.

**Figure 8.**
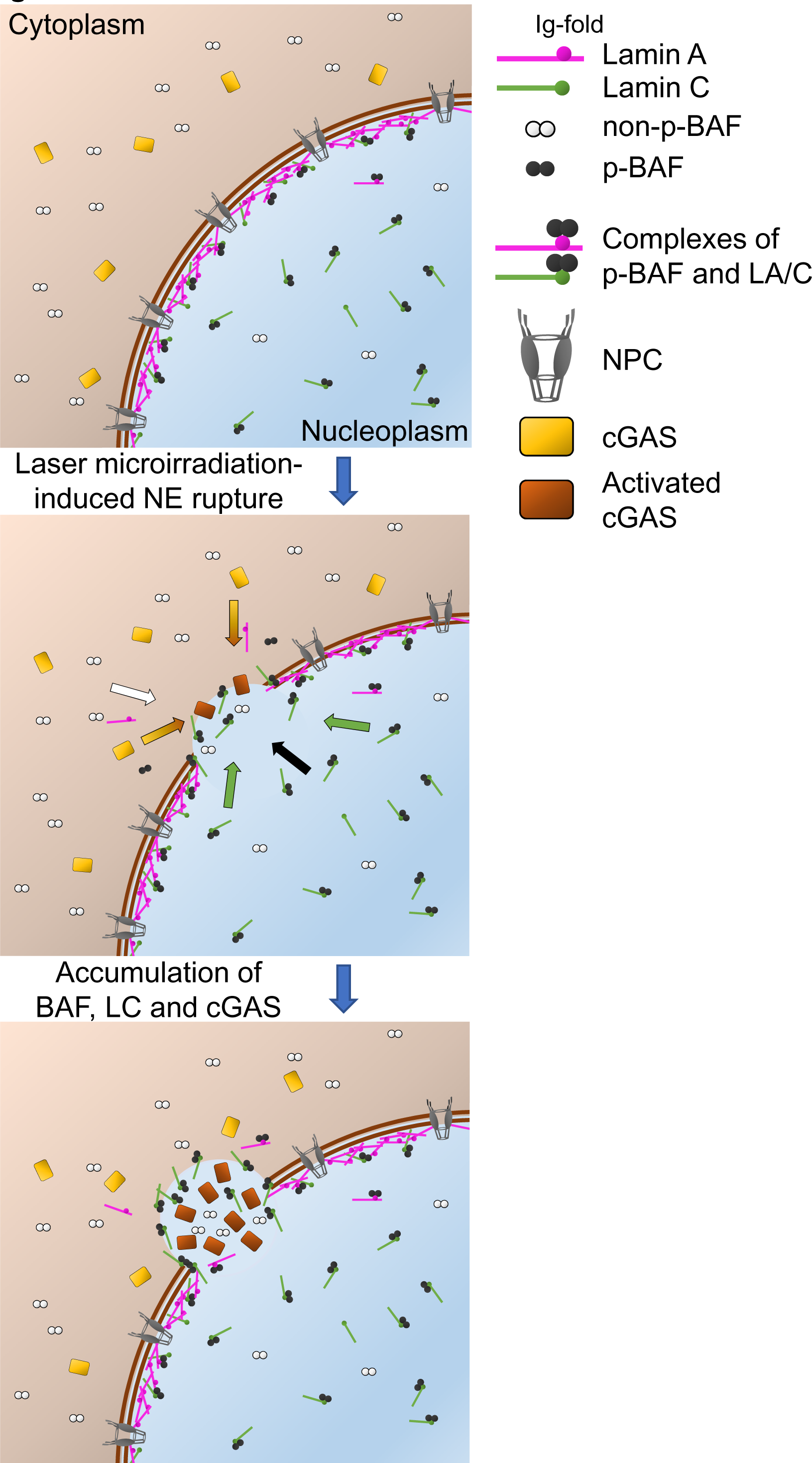
A summary diagram of NE rupture and the repair. NE rupture and the repair are depicted in three sequential steps (before (0 min, top), ∼1 min (middle) and up to ∼10 min (bottom) after laser microirradiation). (Middle) Colored arrows indicate accumulation of non-p-BAF (white), p-BAF (black), LC (green), and cGAS (yellow).

### Differential kinetics of lamin isoforms upon laser-induced NE rupture

Our microscopy analyses demonstrate that LC is clearly the major lamin isoform that accumulates at the rupture sites within 50 s after the rupture. This rapid LC accumulation is explained by its rapid exchange rate in the NL as revealed by FRAP (Broers et al., 1999; Dahl et al., 2006) and the presence of diffusible nucleoplasmic pool (Broers et al., 1999; Shimi et al., 2008). Although LA and LC are expressed at nearly equal levels, the nucleoplasmic pool of LC is much more abundant than that of LA under normal conditions (Kolb et al., 2011; Markiewicz et al., 2002; Wong et al., 2021). Indeed, when overexpressed to increase the nucleoplasmic fraction, mEmerald-LA rapidly accumulates at the ruptured sites. Because the LC-specific six amino-acid sequence at the C-terminus is not required for the rapid accumulation, the extended tail region of LA might facilitate its assembly into filaments and meshwork through the homotypic association (Shimi et al., 2015; Turgay et al., 2017) and/or the interactions with other binding partners such as F-actin (Simon et al., 2010) and SUN1/2 (Crisp et al., 2006; Haque et al., 2006). As LA expression is significantly increased through p53 activation in response to DNA damage (Rahman-Roblick et al., 2007; Yoon et al., 2019), this might contribute to prevent and/or repair of NE ruptures.

### A possible mechanism for the accumulation of LA/C, BAF, and cGAS at the rupture sites

It has been reported that BAF is required for the recruitment of overexpressed GFP-LA to rupture sites (Young et al., 2020). We also show that LC accumulation at rupture sites depends on BAF, supporting the idea that LA/C and p-BAF form diffusible complexes in the nucleus. Recent studies have shown that phosphorylation of both Ser-4 and Thr-3 in BAF by the vaccinia-related kinase 1 (VRK1) greatly reduces the extensive flexibility of the N-terminal helix α1 and loop α1α2 to decrease the affinity for dsDNA but not the Ig-fold domain of LA/C (Marcelot et al., 2021; Nichols et al., 2006; Samson et al., 2018). Therefore, p-BAF in the nucleus might interact transiently with nuclear DNA (Shimi et al., 2004), such that the LA/C-p-BAF complex can freely diffuse throughout the nucleus. In contrast, non-p-BAF strongly binds to dsDNA in the cytoplasm (Kobayashi et al., 2015). Because the phosphorylation-dephosphorylation balance is likely to be altered by the leakage of VRKs (Birendra et al., 2017; Nichols et al., 2006) and the protein phosphatases from the rupture sites, it is tempting to speculate that p-BAF might become dephosphorylated at the rupture sites to restore the strong dsDNA binding affinity. In addition, non-p-BAF can enter the nucleus from the cytoplasm through the opening of the ruptured NE (Halfmann et al., 2019). Although cGAS and BAF can compete for binding to nuclear DNA at the rupture sites (Guey et al., 2020), our data show that cGAS occupies the protruded DNA regions whereas BAF is localized to the peripheries. BAF is associated with the INM through the interaction with LEM domain proteins, which might result in excluding BAF from the DNA regions. Consequently, LA/C could be localized to the peripheries of protruded DNA regions in a BAF-dependent manner.

The BAF-dependent LA/C recruitment mechanism is reminiscent to the mitotic dynamics of LA/C. At the onset of mitosis, LA and LC filaments in the NL are depolymerized into dimers (Gerace and Blobel, 1980; Peter et al., 1990). These dimers are recruited with BAF to the regions of the sister chromosome mass, designated as the ‘core regions’ formed during the anaphase-telophase transition (Haraguchi et al., 2008; Lee et al., 2001). BAF recruits a LEM domain-containing INM protein, LEM2 from the cytoplasm to core regions in mitosis (von Appen et al., 2020), and LEM2 condensates are essential for its accumulation at core regions (von Appen et al., 2020). Because LEM2 is recruited with BAF to the rupture sites (Halfmann et al., 2019), LEM2 may form condensates near the rupture sites to accelerate the accumulation of BAF and LA/C, before LA and LC dimers are polymerized into a filament.

The accumulation of BAF and LC at rupture sites is facilitated when cGAS is present in the cytoplasm. In addition, DNA protrusion becomes significantly evident at the cGAS-positive rupture sites compared to the cGAS-negative sites. The cGAS originated in the cytoplasm might function in DNA decondensation to enlarge the protrusion, which in turn results in increasing the surface area of the INM where BAF and LC are enriched.

Reciprocally to the facilitating function of BAF and cGAS in LA/C accumulation at the rupture sites, the absence of LA/C also affects the BAF and cGAS accumulation. The mechanism of how LA/C contributes to the BAF and cGAS accumulation remains unknown. Emerin might be involved in LA/C-dependent BAF accumulation. GFP-emerin accumulated at the rupture sites in LA/C KD cells (Halfmann et al., 2019), whereas LA/C interacts with emerin through the Ig-fold (Samson et al., 2018), suggesting that emerin might stabilize the LA/C-BAF complexes or condensates underneath the INM at the rupture sites. Because DNA protrusions with cGAS accumulation was dependent on LA/C, it can be speculated that chromatin structure in *Lmna*-KO cells may be altered to reduce the interaction with cGAS.

### Laminopathy-associated mutations causing failure in the rupture site accumulation

In this study, we show that some of the seven laminopathy-associated Ig-fold mutations cause a failure to accumulate at rupture sites. Among them, the five mutants that have weak or no BAF binding activity in vitro do not accumulate at rupture sites whereas the rest two show the decreased levels of the accumulation, even though these mutants show BAF binding activities similar to that of the wild type in vitro. Unlike the diluted conditions at room temperature for in vitro measurements, the crowded nuclear environment at 37°C might provide sensitive detection of the mutation effects. Therefore, the NE rupture assay could be a useful method to functionally analyze laminopathy mutations. Because some of the mutations that are analyzed in this study have been reported to be homozygous, our findings also implicate a possible link between NE rupture and the physiological properties of and pathological changes in the laminopathies.

## Materials and methods

### Plasmid construction

The plasmids used for expressing lamins (mEmerald-LA, mEmerald-LB1, mEmerald-LB2, and mEmerald-LC) were described previously (Shimi et al., 2015). The amino acid substitution mutants were generated by PCR mutagenesis using KOD One PCR Master Mix -Blue-(Toyobo). Primers used in this study are listed in Table 2. LC NLS deletion (Δ417-422) was generated by KOD One PCR amplification and the In-Fusion HD Cloning Kit (Clontech). The NLS of SUN2 (KDSPL**<u>R</u>**TL**<u>KRK</u>**SSNM**<u>KR</u>**L) (Turgay et al., 2010) cDNA was amplified by the PrimeSTAR HS DNA Polymerase (Takara) from pEGFPN3-SUN2 (Nishioka et al., 2016), kindly gifted by Miki Hieda (Ehime Prefectural University of Health Sciences), and used for replacing the NLS of LC to construct the Δ417-422 + NLS^SUN2^ using the In-Fusion. The chimeric LC with the Ig-fold of LB1 (Δ432-548 + Ig-fold^LB1^) was constructed by the In-Fusion with amplified DNA fragments by the KOD One. The pCDH-CMV-MCS-EF1-Blast was generated from pCDH-CMV-MCS-EF1-Hygro (plasmid *#*CD515B-1; System Biosciences). The annealed synthetic oligonucleotides (Table 2, Integrated DNA Technologies) encoding two different NLSs derived from SV40 large T antigen (NLS^SV40^; P**<u>KKKRK</u>**V) and c-Myc (NLS^Myc^; PAA**<u>KR</u>**V**<u>K</u>**LD) (Ray et al., 2015) were ligated to 5’ and 3’ of the sfCherry (Nguyen et al., 2013) using the In-Fusion and the Ligation-Convenience Kit (Nippon Gene), respectively, and designated as pCDH-NLS^SV40^-sfCherry-NLS^Myc^-Blast. The sfGFP cDNA was amplified from sfGFP-C1 (Addgene plasmid # 54579; http://n2t.net/addgene:54579; RRID:Addgene_54579; a gift from Michael Davidson & Geoffrey Waldo) (Pédelacq et al., 2006) using the KOD One, cloned into pCDH-CMV-MCS-EF1-Hygro using the In-Fusion, and designated as pCDH-sfGFP-MCS-EF1-Hygro. The cDNA encoding 6xHis-tagged DARPin-LaA_6 (Zwerger et al., 2015) was ligated into *Not* I/*Eco*R I-digested pCDH-sfGFP-MCS-EF1-Hygro using the Ligation-Convenience Kit and designated as pCDH-sfGFP-DARPin-LA6-Hygro. The pLKO.2 hygro, pLKO.3 blast, pLKO.4 neo, pLKO.5 Zeo, and pLKO.6 sfCherry were generated from pLKO.1 puro plasmid (Addgene plasmid # 8453; http://n2t.net/addgene:8453; RRID:Addgene_8453; a gift from Bob Weinberg) (Stewart et al., 2003). The annealed oligonucleotide encoding scrambled shRNA (shScr, Table 2, Integrated DNA Technologies) was cloned into them using the Ligation-Convenience Kit. For specific KD of LA (Harr et al., 2015) and LC (Wong et al., 2021), shRNA-expressing sequences were cloned into pLKO.2 hygro and pLKO.6 sfCherry using the Ligation-Convenience Kit. The annealed oligonucleotides to express anti-*Banf1*-shRNAs (shBAF#1; TRCN0000124958, and shBAF#2; TRCN0000124955, Table 2, Broad Institute) were also ligated into pLKO.6 sfCherry using the Ligation-Convenience Kit. The cDNA fragment encoding HaloTag from CDK9-Halo (Uchino et al., 2022) was amplified by the KOD One, and the mEmerald sequence of pmEmerald-C1 was replaced by the In-Fusion to construct pHaloTag-C1. The annealed oligonucleotide encoding NLS^SV40^ was also cloned into the N-terminus of HaloTag using the In-Fusion and designated as pNLS^SV40^-Halo. The cDNA encoding mouse BANF1/BAF was synthesized using two120-mers and a 90-mer single-stranded DNA oligonucleotides (Table 2, Sigma-Aldrich). These ssDNA oligonucleotides each contain a 15-mer overlapping sequence at both ends, and were assembled into pHaloTag-C1 to produce the pHalo-Baf in one-step using the Gibson Assembly Master Mix (New England Biolabs). The cGAS cDNA was amplified by the KOD One from pLPC-cGAS-Flag (Dou et al., 2017), kindly gifted by Zhixun Dou (Massachusetts General Hospital and Harvard Medical School), fused to the N-terminus of sfCherry using the In-Fusion, and designated as pcGAS-sfCherry.

**Table 2.**
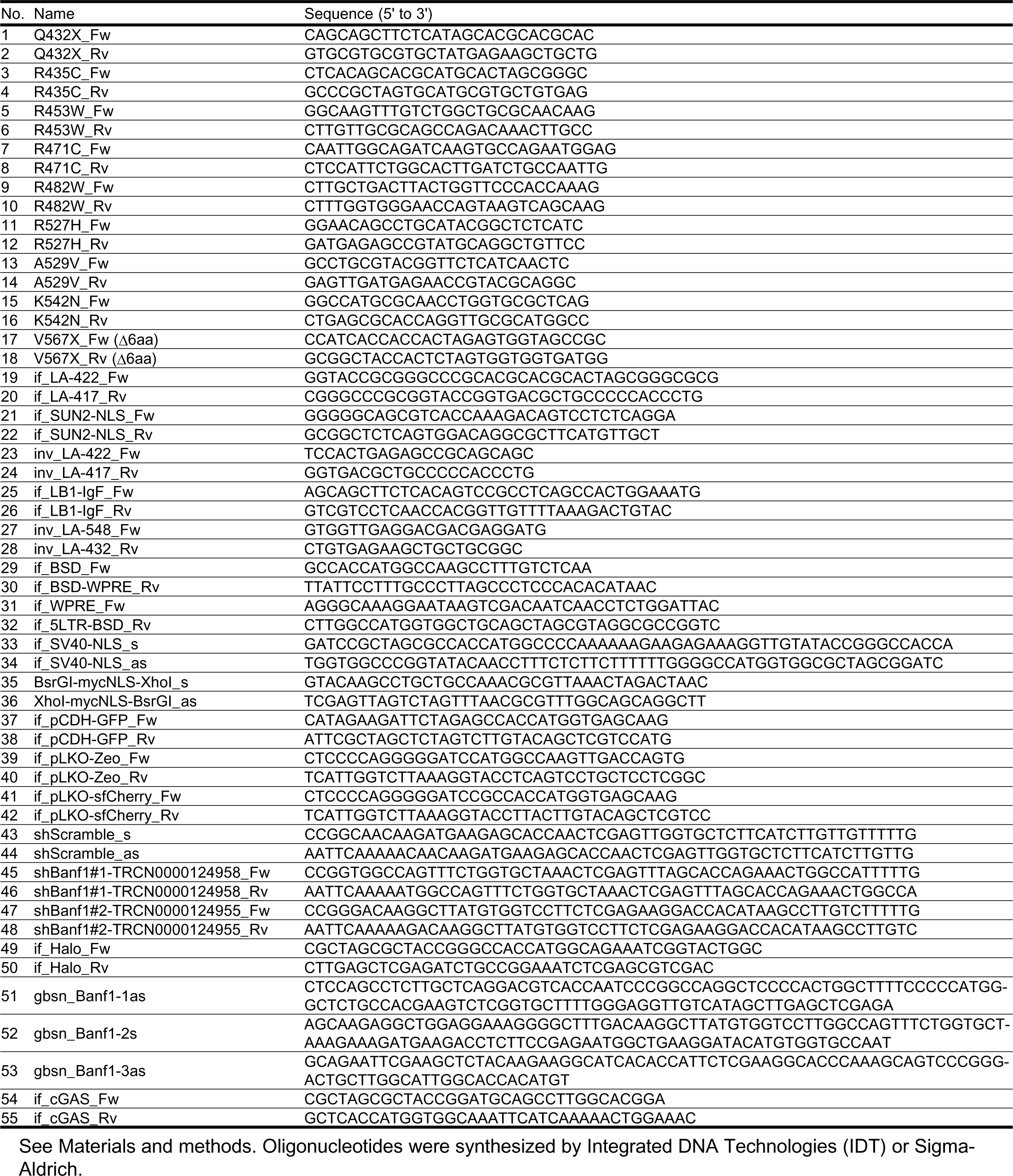
Oligonucleotides used in this study.

### Cell culture and transient transfection

Immortalized WT, *Lmna* ^-/-^, *Lmnb1* ^-/-^, and *Lmnb2* ^-/-^ MEFs (Kim et al., 2011; Kim and Zheng, 2013; Shimi et al., 2015) and C2C12 cells (CRL-1772, ATCC) were cultured in modified DMEM (high glucose; Nacalai Tesque) containing 10% FBS (qualified, Thermo Fisher), 4 mM L-glutamine, 100U/mL penicillin and 100 μg/mL streptomycin (Merck) at 37°C in a humidified chamber with 5% CO2. BJ-5ta cells (CRL-400, ATCC) were cultured in a 4:1 mixture of DMEM and Medium 199 (Thermo Fisher) containing 10% FBS, 4 mM L-glutamine, 100U/mL penicillin and 100 μg/mL streptomycin. MCF10A cells (CRL-10317, ATCC) were cultured in DMEM/F-12, GlutaMAX (Thermo Fisher) supplemented with 5% horse serum (New Zealand origin, Thermo Fisher), 10 μg/mL insulin, 0.5 μg/mL hydrocortisone (FUJIFILM Wako), 100 ng/mL cholera toxin (Merck), 100U/mL penicillin, 100 μg/mL streptomycin and 20 ng/mL recombinant human EGF (AF-100-15, PeproTech).

MEFs were transfected with appropriate plasmids using Lipofectamine 3000 (Thermo Fisher) by reverse transfection. Briefly, the DNA-lipid complex (1.25 μg DNA: 3.75 μL Lipofectamine 3000 in 250 μL Opti-MEM; Thermo Fisher) was added to 1.62×10^5^ cells as they were seeding onto a 35 mm dish with 2 mL of growth medium. Cells transiently expressing mEmerald-lamins, Halo-BAF, and cGAS-sfCherry were observed 48 h after transfection.

### Lentiviral transduction

For lentivirus-mediated stable introduction of sfGFP-DARPin-LA6 and NLS^SV40^-sfCherry-NLS^Myc^, we followed the methods described previously (Liu et al., 2021). Briefly, pVSV-G (PT3343-5, Clontech) and psPAX2 (Addgene plasmid #12260; http://n2t.net/addgene:12260; RRID:Addgene_12260; a gift from Didier Trono), together with the pCDH vector (pCDH-NLS^SV40^-sfCherry-NLS^Myc^-Blast or pCDH-sfGFP-DARPin-LA6-Hygro) in a 1:3:4 weight ratio of each plasmid was transfected into ∼80% confluent 293T cells (CRL-3216, ATCC) using Lipofectamine 3000 following the manufacturer’s instructions for lentivirus production. One day after the transfection, the medium was replaced with fresh medium, which was harvested at 48 h after transfection. For virus infection, MEFs, C2C12, BJ-5ta, and MCF10A cells were incubated with the virus-containing culture supernatants with 4 µg/mL polybrene (Nacalai Tesque) for 24 h. Infected cells were selected by incubation in medium containing 200 µg/mL hygromycin B Gold or 3 µg/mL blasticidin S (InvivoGen) for 4 d, except that C2C12 cells were selected with 20 µg/mL blasticidin S for 4 d.

### Live-cell imaging and NE rupture induction by laser microirradiation

Culture medium was replaced with FluoroBrite (Thermo Fisher) containing 10% FBS, 4 mM L-glutamine, 100U/mL penicillin and 100 μg/mL streptomycin. For imaging Halo-BAF and NLS-Halo, the cells were stained with 0.1 nM Janelia Fluor 646 HaloTag Ligand (Promega) for 30 min. For laser-microirradiation and image collection, cells were set onto a heated stage (37°C, Tokai Hit) with a CO2-control system (Tokken) on a confocal microscope (FV1000; Olympus) with a 60× PlanApo N (NA 1.4) oil lens operated by built-in software FLUOVIEW ver. 4.2 (Olympus). All live-cell images were collected using a main scanner (4% 488-nm laser transmission; 30% 543-nm laser transmission; 0.1% 633-nm laser transmission; 2 μs/pixel; 512×512 pixels; pinhole 100 μm; 6× zoom; ∼ 10 s/frame). After the first image was acquired, a 2 μm diameter spot was laser-microirradiated using a second scanner at 100% power of 405-nm laser transmission (40 μs/pixel, approximately 780 μW) for 10 s while the images were collected with another scanner. The optical power was measured by a laser power meter LP1 (Sanwa) and calculated by the correction coefficient (×8.8 for 405-nm). For photobleach experiments before NE rupture, a 10 μm diameter spot of cell nucleus was photobleached using a main scanner at 100% power of 488-nm laser for 40 s (200 μs/pixel).

Fiji software Image J ver. 1.53k (National Institute of Health) was used for measuring max intensity in the region of interest after Gaussian filtering with σ = 2.0 over time. Fluorescence intensity was normalized by each of the initial intensity in the laser-microirradiated region. For analyzing the accumulation dynamics of sfGFP-DARPin-LA6, cGAS-sfCherry and Halo-BAF, their intensities were normalized by the time point of the peak in each cell, respectively. CellProfiler ver. 3.1.9 (Broad Institute) with the NE_NP_intensity pipeline (programed by Arata Komatsubara, uploaded on GitHub; http://github.com/ArataKomatsubara/NE_NP_intensity) were used for measuring mean intensity of mEmerald-LA/C in the nucleoplasm. Briefly, after the nuclear region was recognized by the nuclear localization of NLS-sfCherry, 3 pixels (207 nm) and more than 10 pixels (690 nm) inside from the rim of nuclear region were regarded as NE and the nucleoplasm, respectively, and then the mean intensity of mEmerald-LA/C was measured.

### Indirect immunofluorescence and microscopy

Primary antibodies used for indirect immunofluorescence were mouse monoclonal anti-LA (1:500; 4A58; sc-71481, or 1:1,000; C-3; sc-518013, Santa Cruz), rabbit polyclonal anti-LA (1:2,000; 323; (Dechat et al., 2007)), rabbit polyclonal anti-LC (1:2,000; 321; (Kittisopikul et al., 2021; Kochin et al., 2014), or 1:200; RaLC; ab125679, Abcam; (Tilli et al., 2003; Venables et al., 2001)), mouse monoclonal anti-LB1 (1:1,000; B-10; sc-5374015 (for mouse cells), or 1:500; 8D1; sc-56144 (for human cells), Santa Cruz), rabbit monoclonal anti-LB2 (1:100; EPR9701(B); ab151735, Abcam), mouse monoclonal anti-NPC proteins (FXFG repeat-containing nucleoporins) (1:1,000; mAb414; 902907, Biolegend), rabbit monoclonal anti-BANF1/BAF (1:200; EPR7668; ab129184, Abcam), mouse monoclonal anti-BANF1 (1:400; 3F10-4G12; H00008815-M01, Abnova) and rabbit monoclonal anti-cGAS (1:250; D3O8O; 31659 (mouse specific), Cell Signaling Technology). The secondary antibodies used were Alexa Fluor 488-donkey anti-mouse immunoglobulin G (IgG), Alexa Fluor 488-donkey anti-rabbit IgG, Alexa Fluor 568-donkey anti-rabbit IgG (1:2,000, 1:2,000 and 1:1,000, respectively, Thermo Fisher), Cy5-donkey anti-mouse IgG and Cy5-donkey anti-rabbit IgG (all 1:2,000, Jackson ImmunoResearch).

Cells were grown on 35 mm Glass Based Dishes or 24-well EZVIEW Glass Bottom Culture Plates LB (IWAKI) and fixed with 4% PFA (Electron Microscopy Sciences) for 15 min, followed by permeabilization using 0.1% Triton X-100 in PBS for 10 min and then blocking with Blocking One-P (Nacalai Tesque) in PBS for 30 min. In case of immunofluorescence after NE rupture, cells were grown on 35 mm Glass Based Dishes which have a cover glass with a grid pattern (IWAKI). Cells were fixed within 10 min after the first laser-microirradiated cell or at 60 min after the last laser-microirradiated with 4% PFA containing 0.1% Triton X-100 (Nacalai Tesque) for 15 min, because it helped to stain NE plaques with primary antibodies after NE rupture. For immunofluorescence of rabbit monoclonal anti-BANF1/BAF (clone EPR7668; ab129184), cells were fixed with 3% PFA for 10 min and then permeabilized using -20°C methanol (Nacalai Tesque) for 10 min followed by 0.4% Triton X-100/PBS for 15 min, in accordance with a previous report (Halfmann et al., 2019). Cells were incubated with primary antibodies overnight at 4°C, washed with 10% Blocking One-P in PBS, and incubated with secondary antibodies for 1 h at RT. DNA was stained with Hoechst 33342 (0.5 μg/mL; Thermo Fisher).

High-resolution fluorescence images of sfGFP-DARPin-LA6, BAF and cGAS were acquired using a confocal microscope (IXplore SpinSR, Olympus), which is equipped with microlenses on the spinning disk, a 3.2× magnification changer and a PlanApoN 60× OSC2 (NA 1.4) oil lens and operated by built-in software cellSens Dimension ver. 3.1 (Olympus). The images were collected using an sCMOS (2048×2048 pixels, ORCA Flash 4, Hamamatsu Photonics) with 20% 405-nm laser transmission, 400 ms; 30% 488-nm laser transmission, 200 ms; 40% 561-nm laser transmission, 200ms; and 20% 640-nm laser transmission, 200 ms. Other confocal fluorescence images were obtained using a confocal microscope FV1000 with a 60× PlanApo N (NA 1.4) oil lens (1.0-14.0% 405-nm laser transmission; 3.1-28.1% 488-nm laser transmission; 0.9-30.0% 543-nm laser transmission; 0.1-9.3% 633-nm laser transmission; 2 μs/pixel; 512×512 pixels; pinhole 100 μm). All images were processed in Photoshop ver. 23 (Adobe) for cropping and brightness/contrast adjustment when applicable.

### Immunoblotting

Primary antibodies used for immunoblotting were mouse monoclonal anti-LA/C (1:20,000; 3A6-4C11, eBioscience), rabbit monoclonal anti-BANF1/BAF (1:500; EPR7668), rabbit monoclonal anti-cGAS (1:1,000; D3O8O) and anti-GAPDH (1:5,000; 6C5, Santa Cruz). The secondary antibodies used were anti-Mouse IgG HRP-Linked Whole Ab Sheep (1:10,000; NA931V, Amersham) and anti-Rabbit IgG HRP-Linked F(ab)^2^ Fragment Donkey (1:10,000; NA9340, Amersham).

Cells were washed with ice-cold PBS twice and lysed with 2× Laemmli sample buffer (4% SDS, 20% glycerol, 120 mM Tris-HCl (pH 6.8)) and then incubated at 95°C for 5 min. The protein concentration was measured using a Protein Assay BCA kit (FUJIFILM Wako) and adjusted to have a same concentration among the samples. After mixing bromophenol blue (at final concentration 0.005%, FUJIFILM Wako) and DTT (at a final concentration 125 mM), cell lysates were further denatured by heating at 95°C for 5 min, separated on polyacrylamide gels (SuperSep Ace, 15% or 7.5%, FUJIFILM Wako), and transferred onto polyvinylidene fluoride membranes (FluoroTrans W, 0.2 μm pore, Pall). After incubation in Blocking One (Nacalai Tesque) at RT for 1 h, the membranes were incubated with the primary antibody in Solution 1 of Can Get Signal (Toyobo) for 1 h with gentle shaking, washed in TBS-T (20 mM Tris-HCl, pH 8.0, 150 mM NaCl, 0.05% Tween 20) three times for 5 min each, and incubated with the secondary antibody in Solution 2 of Can Get Signal for 1 h. After washing with TBS-T three times for 5 min each, the membranes were incubated with ImmunoStar Zeta or LD (FUJIFILM Wako) for 5 min before detecting the signals using a LuminoGraph II Chemiluminescent Imaging System (ATTO). The intensity of bands was measured using CSAnalyzer 4 ver. 2.2.2 (ATTO).

### Generation of the LA-, LC- and BAF-KD MEFs

MEFs expressing shRNAs of anti-LA, anti-LC and anti-BAF were generated through lentiviral transduction. KD by shRNA expression was mediated by Lentivirus transduction (see the section on Lentiviral transduction above). For Fig. 2 E, F, and S2, specific KD of LA or LC was determined by sfCherry expression 4-5 d after lentiviral transduction. For Fig. 4 F and G; S4 B and C, specific KD of BAF was determined by expression of sfCherry 4 d after lentiviral transduction. For Fig. 5 C and D, specific KD of LA and/or LC was carried out by lentiviral transduction and 6 d selection with 100 µg/mL hygromycin B Gold.

### Statistical analyses

Unless mentioned otherwise, all plots showed mean values ± SEM (error bars). The linear mixed model was performed to analyze the interaction between group and time using SPSS Statistics ver. 28 (IBM), where ’subject cell’ was set as a random effect, while group and time as fixed effects. The interaction between group and time (group × time) was also set as a fixed effect. Differences were considered significant if P < 0.05. The Mann-Whitney U test was performed in SPSS Statistics to analyze the leakage of NLS-Halo from the nucleus to the cytoplasm for all time points. The Kruskal-Wallis test followed by the Steel-Dwass post-hoc test was performed to multiple comparisons using EZR on R commander ver. 1.54 (programed by Yoshinobu Kanda) (Kanda, 2013).

### Structural depiction

The structural image of the Ig-fold of human LA/C with colored amino acid residues of laminopathy mutations was generated using a published structure (RCSB Protein Data Bank ID: 1IFR) (Dhe-Paganon et al., 2002) as a template, by the PyMOL Molecular Graphics System ver. 2.5 (Schrödinger).

## Supplemental material

Fig. S1 shows the localization of LA, LC, LB1, and LB2 to the rupture sites in MEFs, and C2C12, BJ-5ta, and MCF10A cells by immunofluorescence. Fig. S2 shows the validation of LA- and LC-KD by immunofluorescence and immunoblotting, and the effect of LC KD on the leakage of NLS-Halo from the nucleus to the cytoplasm by live-cell imaging. Fig. S3 shows dynamics of LC Δ6aa at the rupture sites, and the effect of the mEmerald-LA expression level on the dynamics at the rupture sites by live-cell imaging. Fig. S4 shows dynamics of LC Ig-fold laminopathy mutants at the rupture sites by live-cell imaging, the validation of BAF KD by immunofluorescence and immunoblotting, the effect of BAF-KD on the localization of sfGFP-DARPin-LA6 to a protrusion from the nuclear main body in fixed cells, and the effect of Halo-BAF overexpression on dynamics of LC ΔNLS and ΔTail at the rupture sites by live-cell imaging. Fig S5 shows BAF localization in WT, *Lmna-*, *Lmnb1-*, and *Lmnb2*-KO MEFs by immunofluorescence with different fixation conditions, the accumulation timing of sfGFP-DARPin-LA6, Halo-BAF with cytoplasmic or nuclear cGAS-sfCherry by live-cell imaging, and cGAS localization and expression in WT, *Lmna-*, *Lmnb1-*, and *Lmnb2*-KO MEFs by immunofluorescence and immunoblotting, respectively.

## Acknowledgments

We thank Open Facility Center of Tokyo Institute of Technology for nucleotide sequencing and technical assistance, especially Sachiko Muto for image analysis. We also thank all members of the Kimura lab at Tokyo Institute of Technology, especially Yuko Sato and Harumi Ueno for constructing psfCherry-C1, Misako Tanaka and Akito Ohi for constructing pLKO.2-hygro-shScramble, and Arata Komatsubara for image analysis. This work was supported by JSPS KAKENHI Grant Numbers JP20KK0158 (to T. Shimi, Y. Kono and H. Kimura), JP20K06617 (to T. Shimi), and JP18H05527 and JP21H04764 (to H. Kimura). Y. Kono is an academy scholar at the Specialized Academy for Cell Science (Ohsumi-juku) of the Tokyo Tech Organization for Fundamental Research. The authors declare no competing financial interests.

## Author contributions

T. Shimi conceived the project. Y. Kono performed the experiments and analyzed data. R.D. Goldman produced rabbit polyclonal anti-LA and LC specific antibodies. S.A. Adam provided essential plasmids. Y. Zheng established lamin-KO MEFs. O. Medalia developed anti-LA/C specific DARPin. K.L. Reddy designed anti-LA and LC specific KD shRNAs. Y. Kono and T. Shimi drafted the manuscript. S.A. Adam, R.D. Goldman, and H. Kimura edited the manuscript. T. Shimi, S.A. Adam, H. Kimura and R.D. Goldman supervised the experiments.

## Abbreviations

3D-SIM: three-dimensional structured illumination microscopy
BAF: barrier-to- autointegration factor
C/N: cytoplasmic-to-nuclear intensity
cGAS: cyclic GMP-AMP synthase
cryo-ET: cryo-electron tomography
DARPin: designed ankyrin repeat protein
DCM: dilated cardiomyopathy
ESCRT III: endosomal sorting complex required for transport III
FPLD: familial partial lipodystrophy
HGPS: Hutchinson-Gilford Progeria syndrome
Ig-fold: immunoglobulin-like fold
INM: inner nuclear membrane
KD: knockdown
KO: knockout
LA: lamin A
LB1: lamin B1
LB2: lamin B2
LC: lamin C
LEM: LAP2-emerin-MAN1
LGMD1B: Limb-girdle muscular dystrophy type 1B
LINC: linker of nucleoskeleton and cytoskeleton
MEF: mouse embryonic fibroblast
MD: muscular dystrophy
NE: nuclear envelope
NL: nuclear lamina
non-p-BAF: non-phosphorylated BAF
NPC: nuclear pore complex
ONM: outer nuclear membrane
p- BAF: phosphorylated BAF
sfGFP: superfolder GFP
sfCherry: superfolder Cherry
shRNA: short hairpin RNA
STING: stimulator of interferon genes
VRK1: vaccinia- related kinase 1.

---

**Figure S1.**
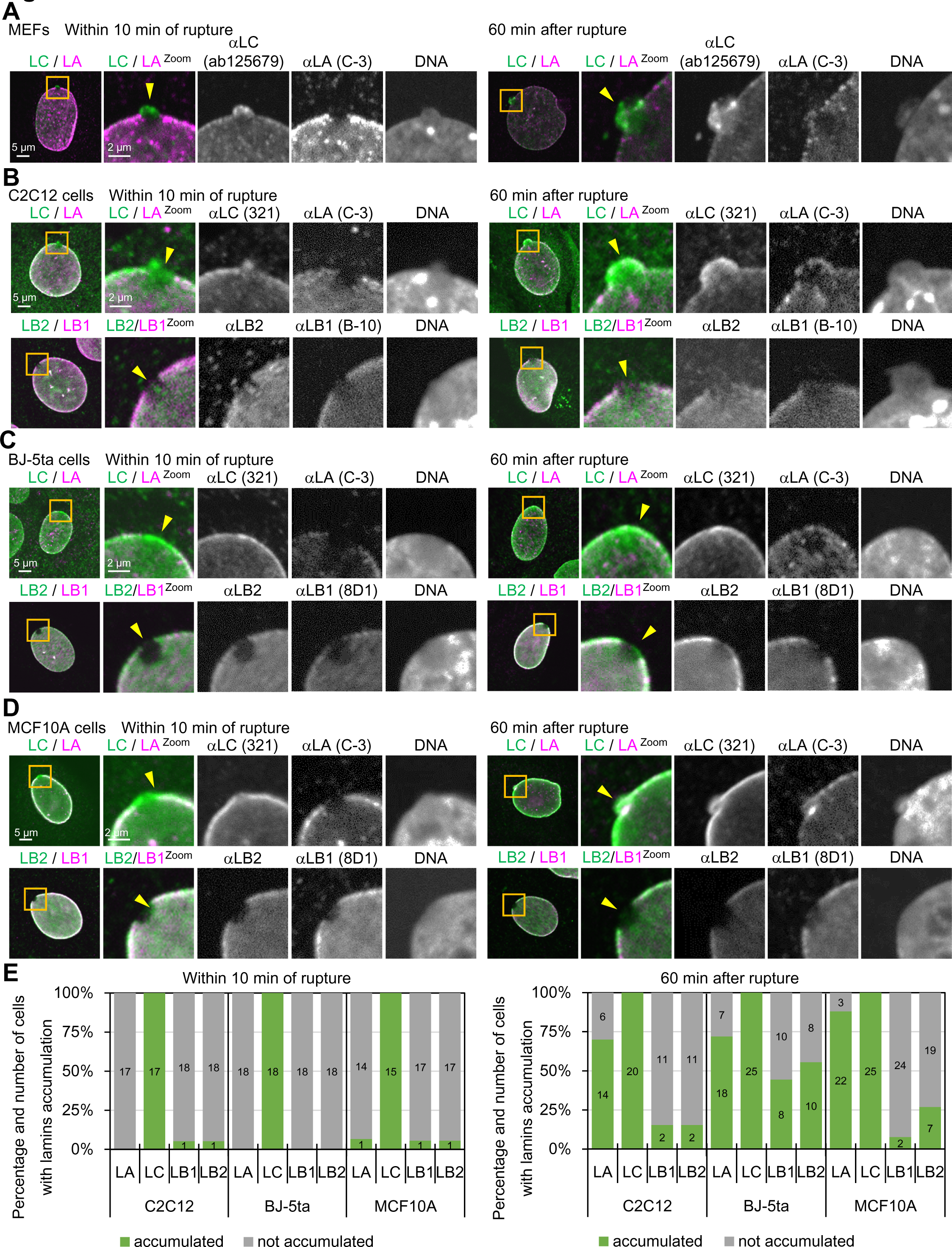
Difference of lamin isoforms in the accumulation kinetics at the rupture sites in MEFs, C2C12, BJ-5ta and MCF10A cells. **(A-D)** A 2-μm diameter spot at the NE in WT MEFs **(A)**, C2C12 **(B)**, BJ-5ta **(C)**, and MCF10A **(D)** were laser-microirradiated to induce the rupture, fixed within 10 min (left panel of each) or 60-70 min later (right panel of each), and then stained with a combination of anti-mouse and anti-rabbit antibodies, followed with Alexa Fluor 488-labeled anti-rabbit IgG and Cy5- labeled anti-mouse IgG, and Hoechst 33342 for DNA. Magnified views of the indicated areas with orange boxes are shown (the second to fifth columns). The ruptured sites are indicated with yellow arrowheads (the second columns). Representative images of single confocal sections. Color-merged images (the first and second columns) in **(A)** MEFs show anti-LC (ab125679, green)/anti-LA (C-3, magenta), **(B)** C2C12 cells show anti-LC (321, green)/anti-LA (C-3, magenta), and anti-LB2 (EPR9701(B), green)/anti-LB1 (B-10, magenta), **(C)** BJ-5ta cells show anti-LC (321, green)/anti-LA (C-3, magenta), and anti- LB2 (EPR9701(B), green)/anti-LB1 (8D1, magenta) and **(D)** MCF10A cells show anti-LC (321, green)/anti-LA (C-3, magenta), and anti-LB2 (EPR9701(B), green)/anti-LB1 (8D1, magenta). Bars: 5 μm (the first column) and 2 μm (the second to fifth column). **(E)** Ratios of cells with (green) and without (gray) enrichments of the indicated antibodies at the rupture sites. The numbers of analyzed cells, fixed within 10 min (left panel) and 60 min (right panel) after laser microirradiation are indicated in the bar charts.

**Figure S2.**
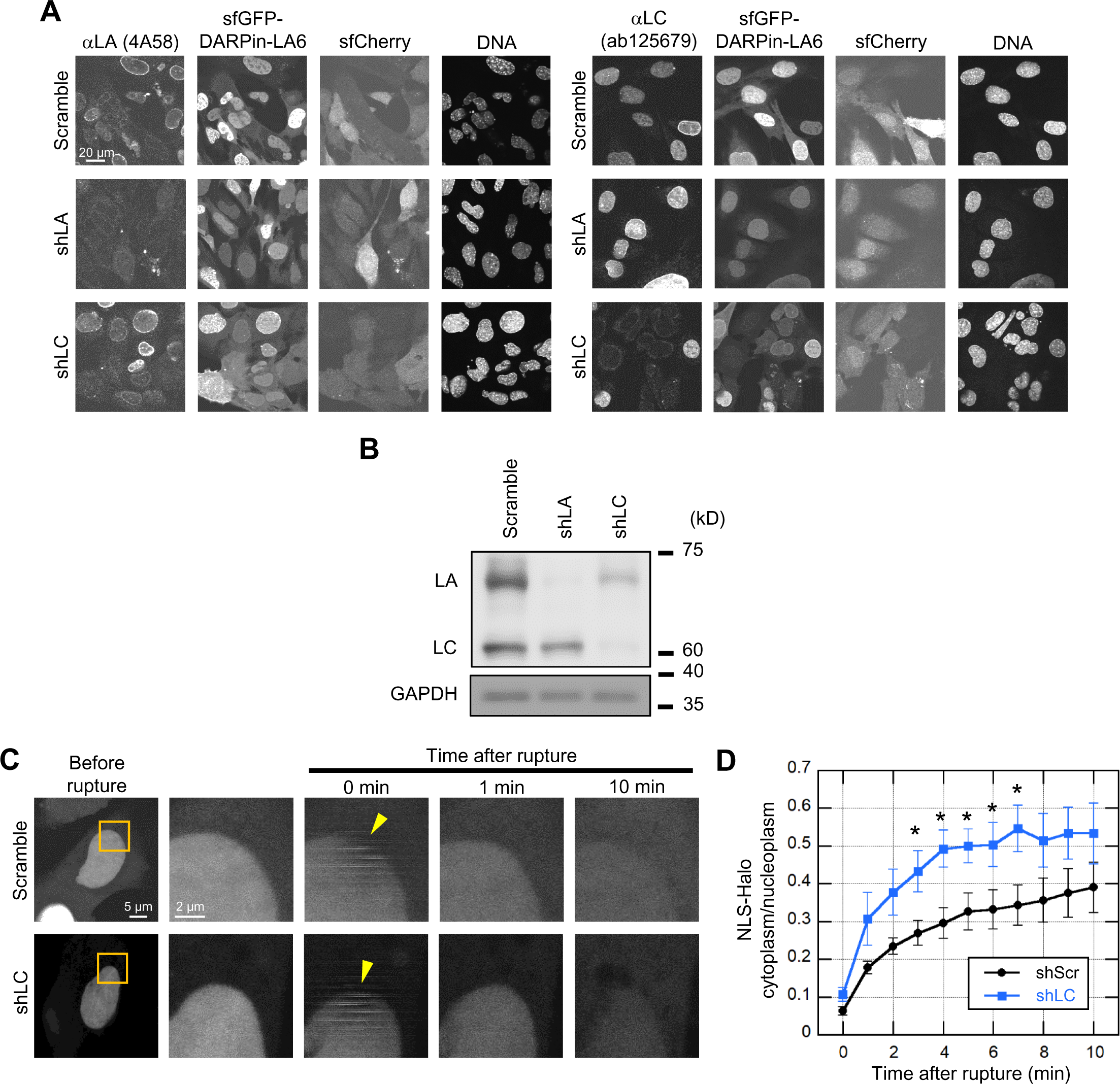
Validation of LA- and LC-KD and the effect of LC KD on the leakage of NLS-Halo from the nucleus to the cytoplasm. **(A and B)** Validation of LA- and LC-KD with immunofluorescence (**A**) and immunoblotting (**B**). **(A)** Representative immunofluorescence images of single confocal sections in WT MEFs expressing scrambled control, shLA or shLC with sfCherry stained with anti-LA (4A58, left panel) and anti-LC (ab125679, right panel), followed with Cy5-labeled anti-mouse and rabbit IgG, respectively, and Hoechst 33342 for DNA. Bar: 20 μm. **(B)** Whole cell lysates from WT MEFs expressing the indicated shRNAs were probed with anti-LA/C and anti-GAPDH (as loading control). Positions of the size standards are shown on the right. **(C** and **D)** During time-lapse imaging of WT MEFs expressing scrambled control (shScr) or shLC with 1 min intervals, a 2-μm diameter spot was laser-microirradiated to induce NE rupture (yellow arrowheads). **(C)** Dynamics of NLS-Halo in response to NE rupture in the indicated cells. **(D)** The cytoplasmic-to-nuclear intensity (C/N) ratio of NLS-Halo was measured and plotted to monitor NE rupture (means ± SEM; *n* = 10 cells; *, P < 0.05 from control by a Mann-Whitney U test).

**Figure S3.**
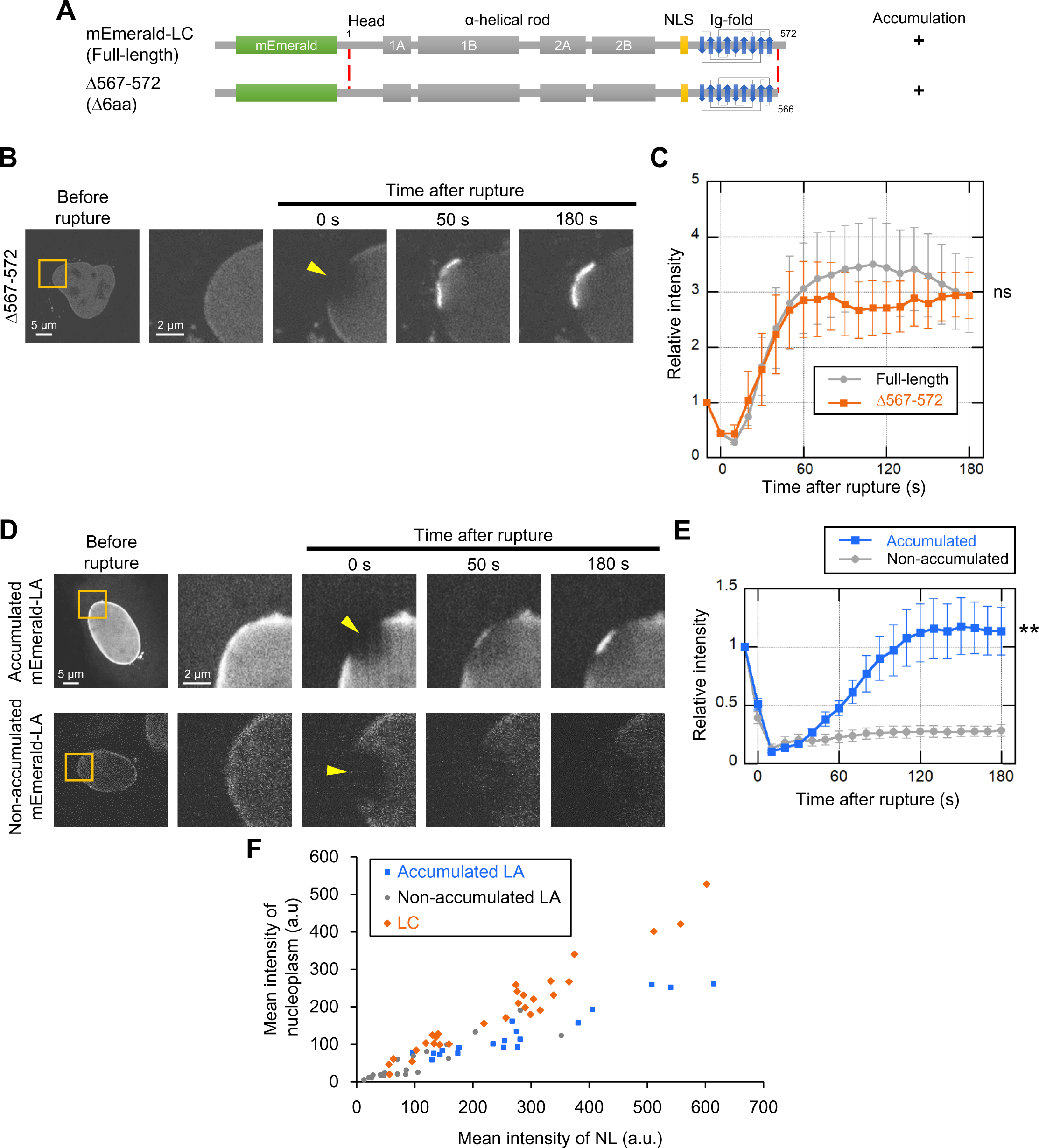
Effects of difference between LA and LC on their accumulation kinetics at the rupture sites. **(A-C)** The requirement of LC-specific 6 amino acids for LC accumulation at the rupture sites. mEmerald-LC full-length and Δ567-572 (Δ6aa) were expressed in *Lmna*-KO MEFs and the NE rupture assay was performed as in Fig 3 A and B. **(A)** Architecture of mEmerald-LC full-length and Δ567-572 (Δ6aa). The summary of their dynamics is indicated on the right (+, accumulated at the rupture sites). **(B)** Dynamics of mEmerald-LC Δ567-572 (Δ6aa) in response to NE rupture. **(C)** Relative fluorescence intensity of the mEmerald-LC Δ567-572 (Δ6aa) (means ± SEM; *n* = 10 cells; ns, P > 0.05 from full-length by a linear mixed model). Full-length (gray) is a reproduction of “Without photobleach” in Fig. 3 B. **(D**-**F)** Relationships between the abundance of nucleoplasmic LA and the accumulation kinetics at the rupture sites. mEmerald-LA were expressed in WT MEFs and the NE rupture assay was performed as in **B** and **C**. **(D)** Dynamics of mEmerald-LA with or without accumulation to the rupture sites. **(E)** Relative fluorescence intensity of the mEmerald-LA (means ± SEM; *n* = 20 cells from two independent experiments; **, P < 0.001 from another by a linear mixed model). Non-accumulated (gray) is a reproduction of “mEmerald-LA” in Fig. 2 B. **(F)** Fluorescence intensities of the mEmerald-LA and LC in the nucleoplasm and the NL was measured before laser microirradiation. (**B** and **D**) The right four columns are magnified views of orange boxes, and the rupture sites are indicated with yellow arrowheads. Bars: 5 μm (the first column) and 2 μm (the second to fifth column).

**Figure S4.**
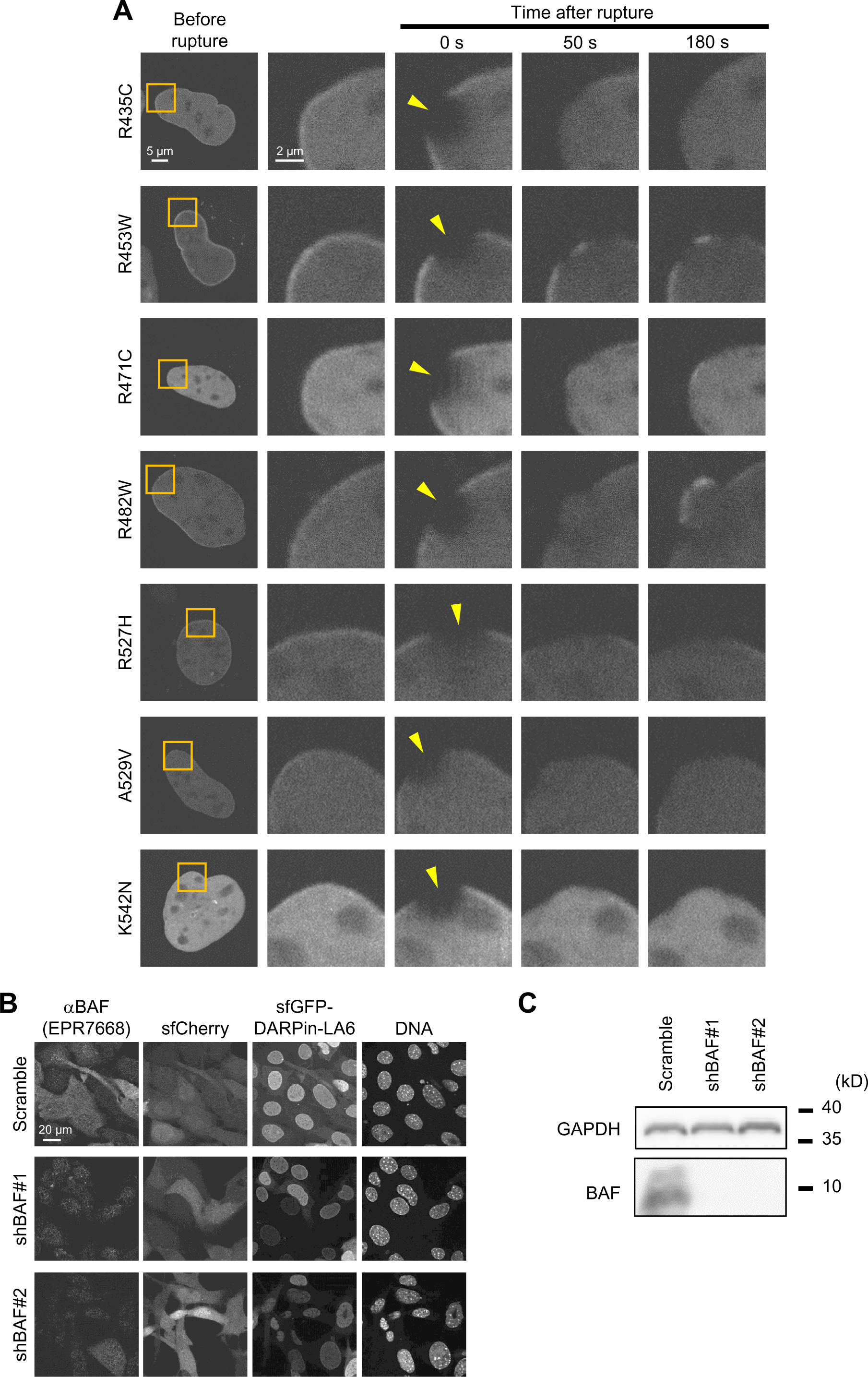
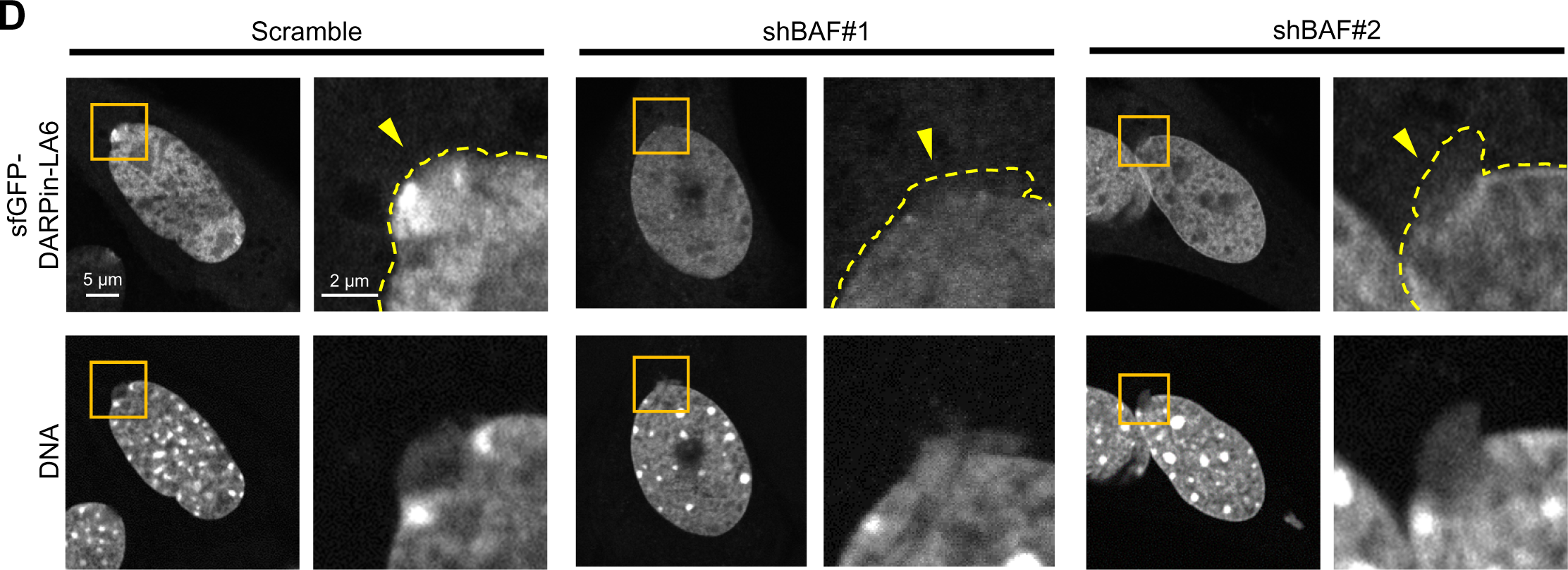
Dynamics of LC Ig-fold laminopathy mutants, validation of BAF KD, and the effect of BAF overexpression on accumulation kinetics of the LC mutants at the rupture sites. **(A)** Dynamics of mEmerald-LC Ig-fold point mutants in *Lmna*-KO MEF. The right four columns are magnified views of orange boxes, and the rupture sites are indicated with yellow arrowheads. Bars: 5 μm (the first column) and 2 μm (the second to fifth column). **(B and C)** Validation of BAF-KD with immunofluorescence **(A)** and immunoblotting (B). **(B)** Representative immunofluorescence images of single confocal sections in WT MEFs expressing scrambled control (shScr), shBAF#1 or shBAF#2 with sfGFP-DARPin-LA6 and sfCherry stained with anti-BANF1/BAF (EPR7668), followed by Cy5-labeled anti-rabbit IgG, and Hoechst 33342 for DNA. Bar: 20 μm. **(C)** Whole cell lysates from MEFs expressing the indicated shRNAs were probed with anti- BANF1/BAF (EPR7668) and anti-GAPDH (as loading control). Positions of the size standards are shown on the right. **(D)** Representative images of single confocal sections of sfGFP-DARPin-LA6 in a NE protrusion in MEFs fixed within 10 min after microirradiation. DNA was stained with Hoechst 33342. The right image of each column is magnified view of orange box. The edges of protruded DNA regions are indicated with yellow dotted line (top right of each column). **(A and D)** Bars: 5 μm (the first column) and 2 μm (the second column to others).

**Figure S5.**
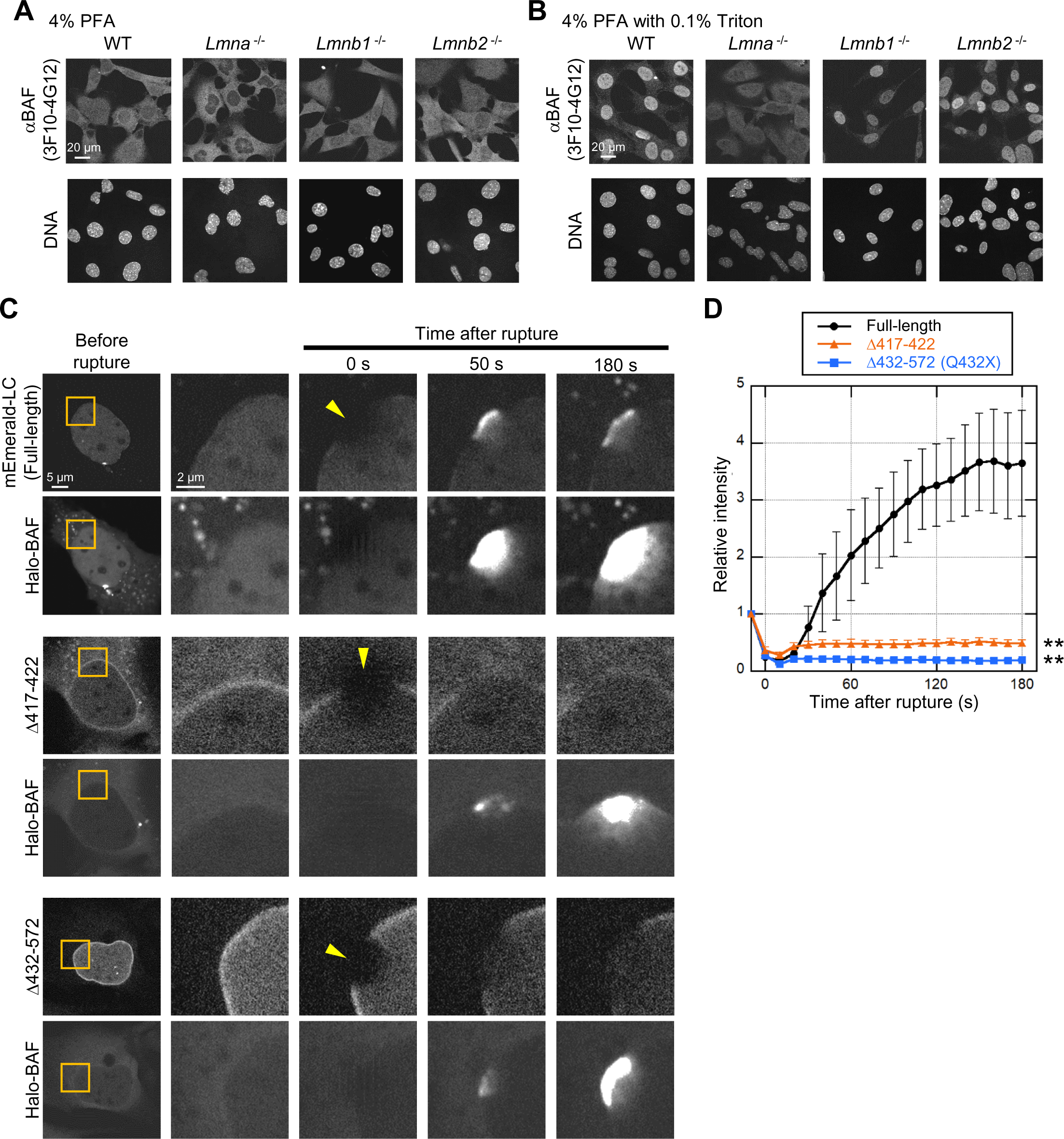
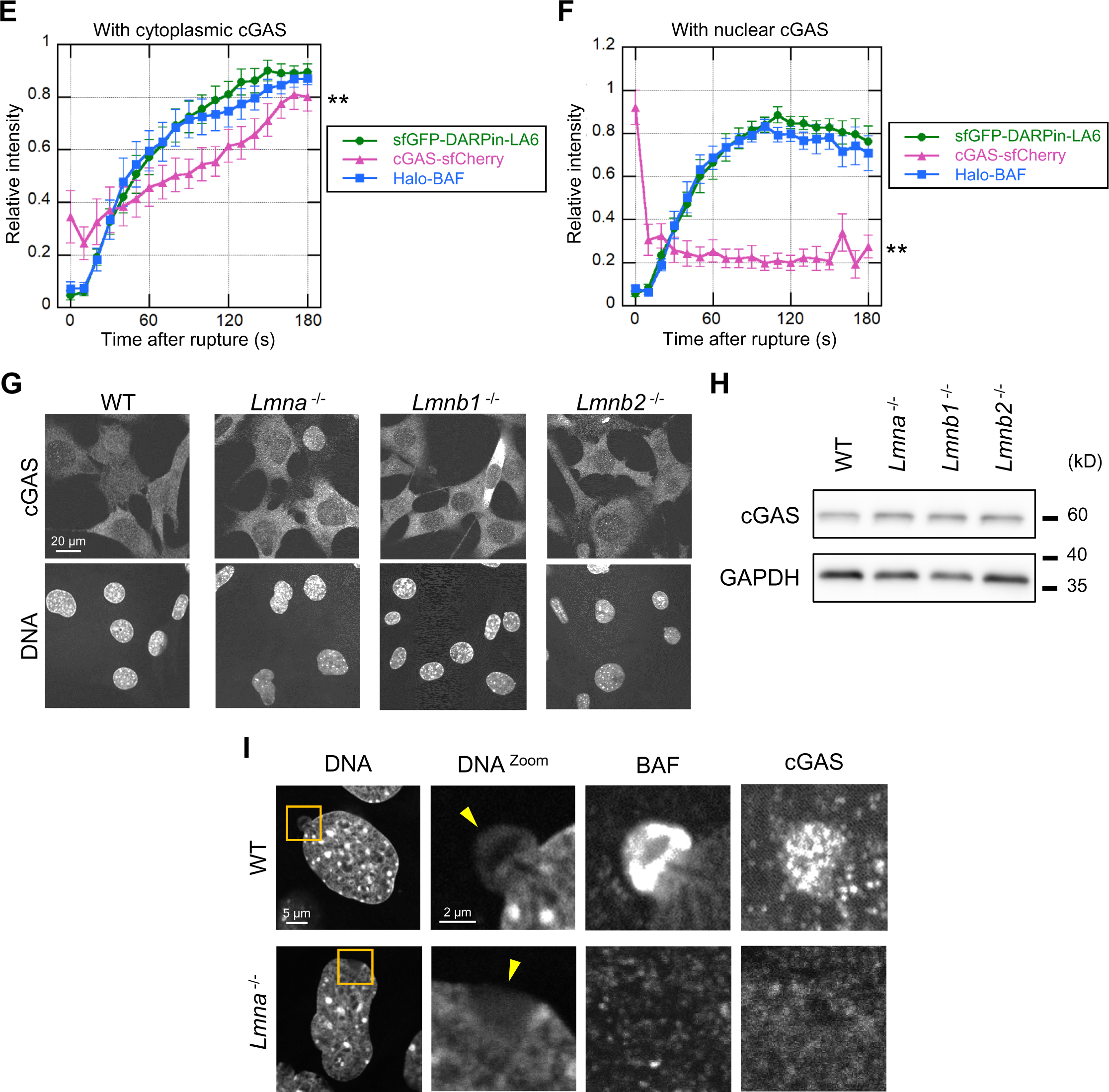
Validation of specific antibodies against BAF and cGAS in lamin-KO MEFs, and the accumulation kinetics of LC, BAF and cGAS at the rupture sites. **(A and B)** Validation of anti-BANF1 (3F10-4G12) for immunofluorescence after different fixation methods. Representative immunofluorescence images of single confocal sections from WT, *Lmna* ^-/-^, *Lmnb1* ^-/-^, and *Lmnb2* ^-/-^ MEFs fixed with 4% PFA only **(A)** or 4% PFA containing 0.1% Triton X-100 **(B)** and stained with the anti-BANF1, followed with Cy5- labeled anti-mouse IgG, and Hoechst 33342 for DNA. **(C** and **D)** The effect of BAF overexpression on accumulation kinetics of the LC mutants at the rupture sites. Halo-BAF (lower panels) with mEmerald-LC full-length, Δ417-422 (ΔNLS) and Δ432-572 (ΔTail) (all, upper panels) were expressed in *Lmna*-KO MEFs. **(C)** Dynamics of mEmerald-LC full-length, Δ417-422 (ΔNLS), Δ432-572 (ΔTail), and Halo-BAF. The right four columns are magnified views of orange boxes, and the rupture sites are indicated with yellow arrowheads. **(D)** Relative fluorescence intensities of the mEmerald-LC mutants (means ± SEM; *n* = 10 cells; **, P < 0.001 from full-length by a linear mixed model). **(E** and **F)** Normalized fluorescence intensities of sfGFP-DARPin-LA6, cGAS- sfCherry and Halo-BAF in MEFs are shown up to the signal peaks, and cGAS-sfCherry is localized to the cytoplasm (**E**) or the nucleus (**F**) (means ± SEM; *n* = 10 cells; **, P < 0.1 from others by a linear mixed model). **(G)** Representative immunofluorescence images of single confocal sections from the indicated MEFs stained with anti-cGAS, followed with Alexa 488-labeled anti-rabbit IgG, and Hoechst 33342 for DNA. **(H)** Whole cell lysates from the indicated MEFs were probed with anti-cGAS, and anti-GAPDH (as loading control). **(I)** Representative immunofluorescence images of single confocal sections from the indicated MEFs fixed within 10 min after laser microirradiation and stained with anti-BANF1 (3F10-4G12) and anti-cGAS, followed with Alexa Fluor 488-labeled anti-rabbit IgG, Cy5-labeled anti-mouse IgG, and Hoechst 33342 for DNA. The right four columns are magnified views of orange boxes, and the rupture sites are indicated with yellow arrowheads. (**A**, **B** and **G**) Bars: 20 μm. (**C** and **I**) Bars: 5 μm (the first column) and 2 μm (the second to others). (**H**) Positions of the size standards are shown on the right.

